# *Nucleocytoviricota* viral factories are transient organelles made by liquid-liquid phase separation

**DOI:** 10.1101/2024.09.01.610734

**Authors:** Sofia Rigou, Alain Schmitt, Anaísa B Moreno, Audrey Lartigue, Lucile Danner, Claire Giry, Feres Trabelsi, Lucid Belmudes, Natalia Olivero-Deibe, Hugo Le Guenno, Yohann Couté, Mabel Berois, Matthieu Legendre, Sandra Jeudy, Chantal Abergel, Hugo Bisio

## Abstract

Phase separation is a widespread mechanism in viral processes, mediating replication, host manipulation and virion morphogenesis. The phylum *Nucleocytoviricota* encompasses diverse and ubiquitous viruses, including *Poxviridae,* the climate-modulating *Emiliania huxleyi* virus and other so-called Giant Viruses. Cytoplasmic members of this phylum form viral factories but their nature has remained unresolved. Here, we demonstrate that these viral factories are formed by liquid-liquid phase separation. We prove that mimivirus viral factories are formed by multilayered phase separation, orchestrated by at least two scaffold proteins. To extend these findings across the phylum *Nucleocytoviricota*, we developed a bioinformatic pipeline to predict scaffold proteins based on a conserved molecular grammar, despite major primary sequence variability. Scaffold candidates were validated in *Marseilleviridae* and *Poxviridae*, highlighting a role of H5 as a scaffold protein in the vaccinia virus. Finally, we provide a repertoire of client proteins of the nucleus-like viral factory of mimivirus and demonstrate important sub-compartmentalization of functions, including those related to the central dogma. Overall, we reveal a new mechanism for an organelle to deploy nuclear-like functions entirely based on phase separation and re-classified phylum *Nucleocytoviricota* viral factories as biomolecular condensates.

## Introduction

Multiple viruses generate biomolecular condensates, which are formed by phase separation (PS) and allow viral replication, host manipulation and virion morphogenesis^1,2^. PS is the process by which biomolecules, such as proteins and nucleic acids, demix into distinct, membrane-less compartments^1–4^. Two types of proteins are concentrated in these compartments: scaffold proteins, which drive the formation and maintenance of these phases, and client proteins, which are selectively recruited to perform specific functions^1–4^. The viral factories (VFs) of several viruses (DNA and RNA) are formed by PS, including members of herpesvirus (with some degree of controversy^5^), adenovirus and reovirus families, as well as of the order *Mononegavirales* (reviewed in^1^). Members of the phylum *Nucleocytoviricota* (including *Poxviridae* and the so-called Giant Viruses^6,7^) generate VFs, but the molecular mechanism behind their biogenesis is currently unknown^8^.

Here, we took advantage of the newly developed genetic tools for giant viruses to study the nature of the VFs of the phylum *Nucleocytoviricota*^9–12^. Particularly, we focus on the VF of mimivirus since it displays a biphasic nature when imaged by electron microscopy (Figure 1A, Figure S1) and a highly synchronized biogenesis and maturation^8^. Moreover, mimivirus is a well-studied member of the family *Mimiviridae,* which are ubiquitous in the environment^13,14^. Mimivirus is a fully cytoplasmic giant virus for which virions are internalized into the host cell by phagocytosis^15^. The acidic compartment of the phagosome, in combination with oxidative stress through Fenton reactions, triggers the opening of the stargate structure located at an apex of the icosahedral capsid^16–18^. Once opened, the fusion of the membrane of the phagosome with the virion’s internal membrane leads to the release in the cytoplasm of the core^15,19,20^. The core contains the genome of the virus and the machinery needed to establish infection^21^. The core consequently develops to establish a viral factory, a highly dynamic and complex organelle-like structure putatively formed by over 300 proteins^22^. VF not only protects the viral DNA from cellular insults^11^ but also segregates DNA replication and transcription from translation^22^.

**Figure 1.**
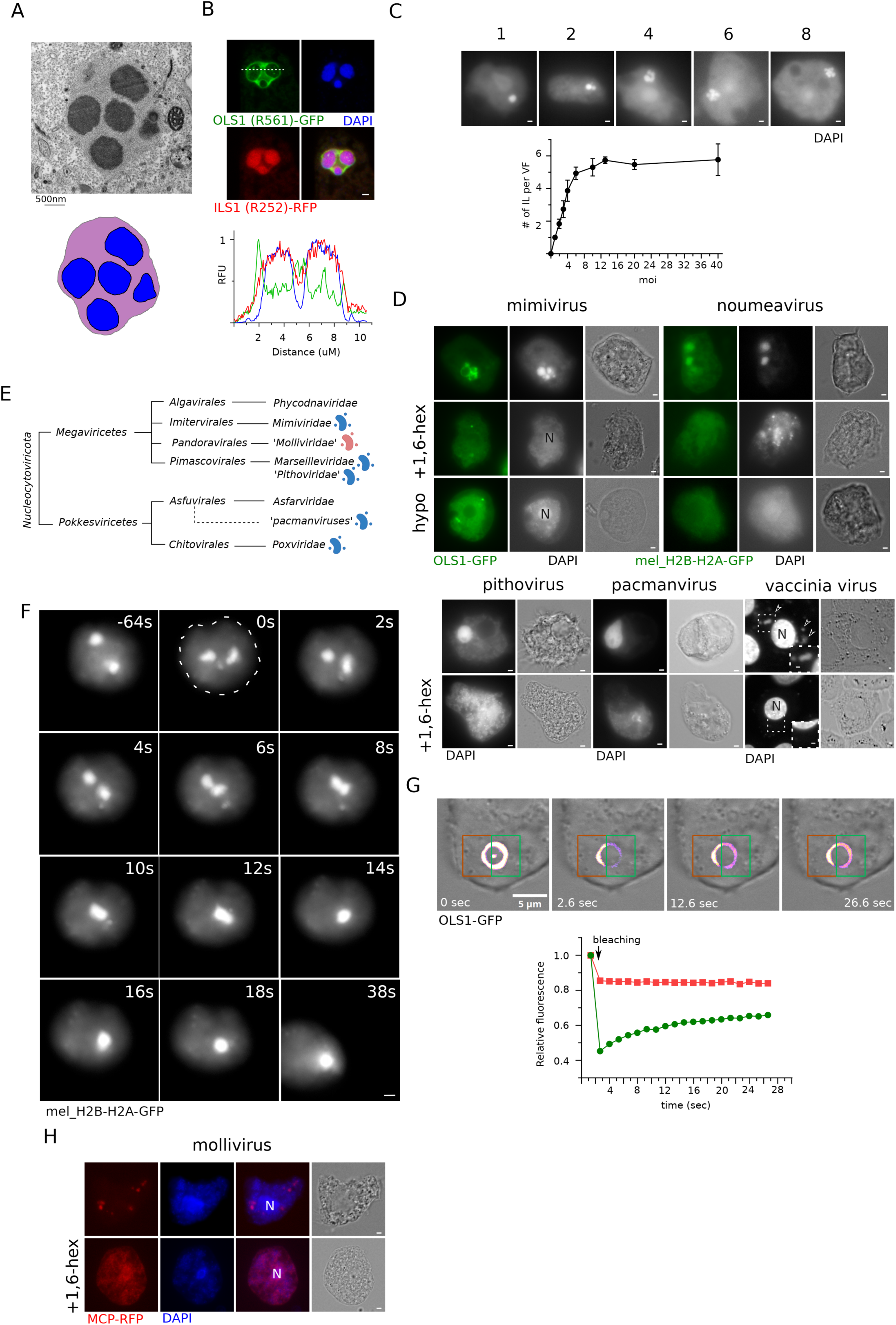
*Nucleocytoviricota* viral factories (VFs) are biomolecular condensates. (A) Transmission electron microscopy imaging of an ultrathin section of infected *A. castellanii* cell with mimivirus VF formed in the cytoplasm. Image was acquired 6h post infection (pi) at a MOI=20. Scale bar: 500nm. A cartoon representing the two layers of the viral factory is also shown. Inner layer (IL) is shown in blue while outer layer (OL) is shown in purple. (B) *A. castellanii* cells expressing C-terminally tagged OLS1-GFP (R561) and ILS1-RFP (R252) were infected with mimivirus. OLS1-GFP localized to the OL of the VF and ILS1-RFP to the IL of the VF. DAPI: DNA. Scale bar: 1μm. Line profiles (right) corresponding to the white dashed line show fluorescence patterns. Image was acquired 6hpi at a MOI=10. (C) Representative light fluorescence microscopy images of *A. castellanii* cells infected with mimivirus harboring VF with different numbers of ILs. Quantification of the number of ILs present in a VF in function of the MOI is shown below the fluorescence images. Data correspond to the mean ± SD of 3 independent experiments. Scale bar: 1μm. (D) Representative light fluorescence microscopy images of *A. castellanii* cells infected with different viruses belonging to *Nucleocytoviricota*. VFs were labelled using DAPI (to label all viral factories) and ectopically expressed OLS1-GFP or mel_H2B-H2A-GFP to label the viral factories of mimivirus or noumeavirus, respectively. Treatment with 10% 1,6-hexanediol or hypotonic media (5% PYG in water) was performed for 10 or 15 minutes (respectively) after mimivirus (3 hpi), noumeavirus (2 hpi), Pithovirus (6hpi) and pacmanvirus (6hpi) infection. Scale bar: 1μm. Representative light fluorescence microscopy images of Vero cells infected with vaccinia virus. VFs were labelled using DAPI and treatment with 10% 1,6-hexanediol was performed for ten minutes after 2hpi. Scale bar: 1μm. Unfilled arrowhead indicate the VFs while the nucleus of the host cell is highlighted with “N”. A zoom of the perinuclear zone is shown in the inset. (E) Taxonomy of viruses belonging to the phylum *Nucleocytoviricota*. Families with a member where 1,6-hexanediol dissolved their VF (blue) or the proposed “Virion Factory” (red) are indicated. We define “Virion Factory” as a phase separated compartment that is not used for genome replication but aids in particle/virion morphogenesis. Image was adapted from ^7^. *Pandoravirales* and probably *Algavirales* englobe viruses with a nucleocytoplasmic infectious cycle (genome is replicated in the nucleus of the host while particles are synthesized at the cytoplasm). (F) Live-cell imaging of noumeavirus infected-*A. castellanii* expressing mel_H2B-H2A-GFP as a marker of the VF. Infection was allowed to proceed for 2h and recording was performed. A single cell is shown and the cell boundaries are indicated by a dashed line in the frame at time 0s. Scale bar: 2μm. (G) Acanthamoeba cells expressing OLS1-GFP were infected with mimivirus at and MOI of 5. Five hours post infection, OLS1-GFP in living cells was photobleached and fluorescence recovery was imaged by confocal microscopy. FRAP data were normalized corrected for background and represented in the recovery curve. (H) Representative light fluorescence microscopy images of *A. castellanii* cells expressing C-terminally RFP-tagged MCP. Representative images of cells infected with mollivirus are shown and treatment with 10% 1,6-hexanediol was performed for ten minutes 6h pi. Scale bar: 1μm. Mollivirus genome replicates at the host nucleus. N: nucleus.

Here, we provide evidence that virtually all VFs produced by members of the phylum *Nucleocytoviricota* are generated by liquid-liquid PS and predict putative scaffold proteins in other members discovered so far. Moreover, we characterize the VF of mimivirus demonstrating a biphasic behavior accomplished by at least 2 scaffold proteins. Client proteins of the VF of mimivirus were identified and their sub-compartmentalization dissected. Overall, we classified VFs of cytoplasmic viruses of the phylum *Nucleocytoviricota* as biomolecular condensates and demonstrated that PS supports nuclear-like functions of mimivirus VF.

## Results

### *Nucleocytoviricota* viral factories are biomolecular condensates

While mimivirus VFs successfully seclude DNA and putatively over 300 proteins^22^, distinct delimiting structures are lacking when imaged by electron microscopy (Figure 1A). Moreover, previous reports indicated that the mimivirus infection starts with the formation of “transport vesicles” (interpreted by us as condensates) that coalesce to form the VFs^8^. Coalescent events of the electron-dense inner layer (IL) of the VF can also be detected when superinfected cells are imaged by EM (Figure S1A). These observations suggest that a PS phenomenon is involved in mimivirus VF formation.

To gather information associated with the fine ultrastructure of VFs, we first fluorescently labelled proteins enriched in a previous proteome of mimivirus VFs^22^ (described later in the manuscript) and identified two clear sub-compartments (Figure 1B). Multiple ILs can be seen in a single VF early during infection, while a single and continuous outer layer (OL) seems to englobe all ILs (Figure 1A-B and Figure S1A). Moreover, the number of ILs per VF linearly increased with the multiplicity of infection (MOI) up to an MOI of approximately 6 (when it becomes difficult to count precisely the number of ILs). These results strongly suggest that each genome unit delivered into the cytoplasm of the host (enclosed in the so-called “core”) has direct repercussions on the number of ILs (Figure 1C). Then, we addressed whether this membrane-less organelle possesses a liquid-like nature by treating infected cells with 10% 1,6-hexanediol (an organic compound proven to interfere with condensate formation due to its ability to alter hydrophobic interactions) for 10 minutes^2^. Mimivirus VF, DNA and protein, partially dissolved during the treatment (Figure 1D and Figure S1B). Treatment with 1,6-hexanediol did not affect the infected cells overall DNA or protein content (Figure S1C-D), demonstrating that the observed changes (Figure 1D and Figure S1B) are due to localization differences. Moreover, treatment with 10% 1,6-hexanediol also dissolved the VFs of noumeavirus (*Marseilleviridae*), pithovirus, pacmanvirus and vaccinia virus (*Poxviridae*) (Figure 1D and Figure S1B), suggesting a shared mechanism for VF formation across the cytoplasmic viruses in the phylum *Nucleocytoviricota* (Fig 1E). 1,6-hexanediol treatment also affected the localization of a VF-localized histone protein from *Marseilleviridae*^12^ (Figure 1D) and did not change the overall DNA or protein content of noumeavirus infected cells (Figure S1C-D). Importantly, 1,6-hexanediol has several known off-target effects, including components of the cytoskeleton^23^ and the inhibition of kinases and phosphatases^24^, all of which could affect VFs independently of liquid-liquid PS. Thus, we subjected infected cells to hypotonic stress (Figure 1D and Fig S1E-F). Hypotonic stress induces swelling of the cells, effectively reducing the concentration of proteins in them^25^. Since PS depends on scaffold protein concentrations, dissolution of the compartment is expected if concentrations are sufficiently low. VFs of mimivirus, noumeavirus, pithovirus and pacmanvirus were also partially dissolved upon hypotonic stress (Figure 1D and Figure S1E-F), reinforcing a liquid-liquid PS behavior of the organelles.

Live-cell imaging is usually used to demonstrate a liquid-like behavior of a biomolecular condensate by showing coalescence and fluorescence recovery after photobleaching (FRAP)^26^. Imaging of a mimivirus infected cell revealed that upon contact, two mimivirus VFs juxtaposed and never separated during the remaining recording time (Figure S1G), suggesting that the OL of two independent VFs fused as observed by fluorescence and electron microscopy (Figure 1A-B). Regardless, these experiments are challenging due to the high mobility of the amoeba and the infrequency of individual mimivirus viral factories (independent ILs without a shared OL) within the same cell. Moreover, relaxation to spherical structures could not be seen, likely due to the presence of the IL. Thus, to simplify the system, we labeled noumeavirus’ VFs, which contain a single detectable layer; multiple VFs can be easily observed early in infection, and noumeavirus infection rapidly disrupts the movement of the amoeba. Cells expressing the C-terminal GFP-tagged mel_H2B-H2A^12^ were infected with noumeavirus and subjected to live-imaging (Figure 1F). Noumeavirus VFs were observed as dynamic organelles subjected to deformation, and the encounter of two VFs yielded a fusion product, showing coalescence. Last, to further reinforce the fluid nature of the VFs of mimivirus, we performed FRAP experiments (Figure 1G). Rapid recovery of fluorescence was observed when an area of the OL of the VF was photobleached (Figure 1G). Moreover, the fluorescence gain from the bleached side was accompanied by a decrease in fluorescence from the other side, demonstrating the diffusion of fluorescent molecules (Figure 1G).

A subgroup of members of the phylum *Nucleocytoviricota* transfer their genome to their host nucleus and replicate their genome there. Mollivirus is one of those viruses^27^. We have previously shown that mollivirus major capsid protein (virion protein) accumulates in an uncharacterized sub-compartment in infected cells before being incorporated into the viral particles^11^. This localization could also be disrupted by 1,6-hexanediol treatment, indicating that particle formation at the cytoplasm of infected cells might also involve PS (Fig 1H and Fig S1B).

Taken together, we concluded that the VFs of mimivirus, noumeavirus, and probably all members of the phylum *Nucleocytoviricota* display characteristics of biomolecular condensates made by liquid-liquid PS.

### At least two scaffold proteins play key roles in mimivirus viral factories’ phase separation

PS is driven by the multivalency of proteins termed scaffold proteins^28^. Scaffolding proteins are abundant in biomolecular condensates and tend to contain intrinsically disordered regions (IDRs), which are compacted means to achieve multivalency^28^. To identify proteins involved in PS in mimivirus, the over 300 viral proteins present in purified VFs^22^ were analyzed for the presence of intrinsically disordered regions using flDPnn^29^. Among these, 43 proteins were found to contain IDRs. Of these, two proteins (Figure 2A) were consistently enriched in three independent immunoprecipitations of VF proteins (baits: R562, R505 and R336/R337), using formaldehyde as a crosslinker agent to preserve weak interactions associated with liquid-liquid PS. These results indicate that these two prey proteins interact with all three client proteins (Supplementary Table 1 and Figure S1H).

**Figure 2.**
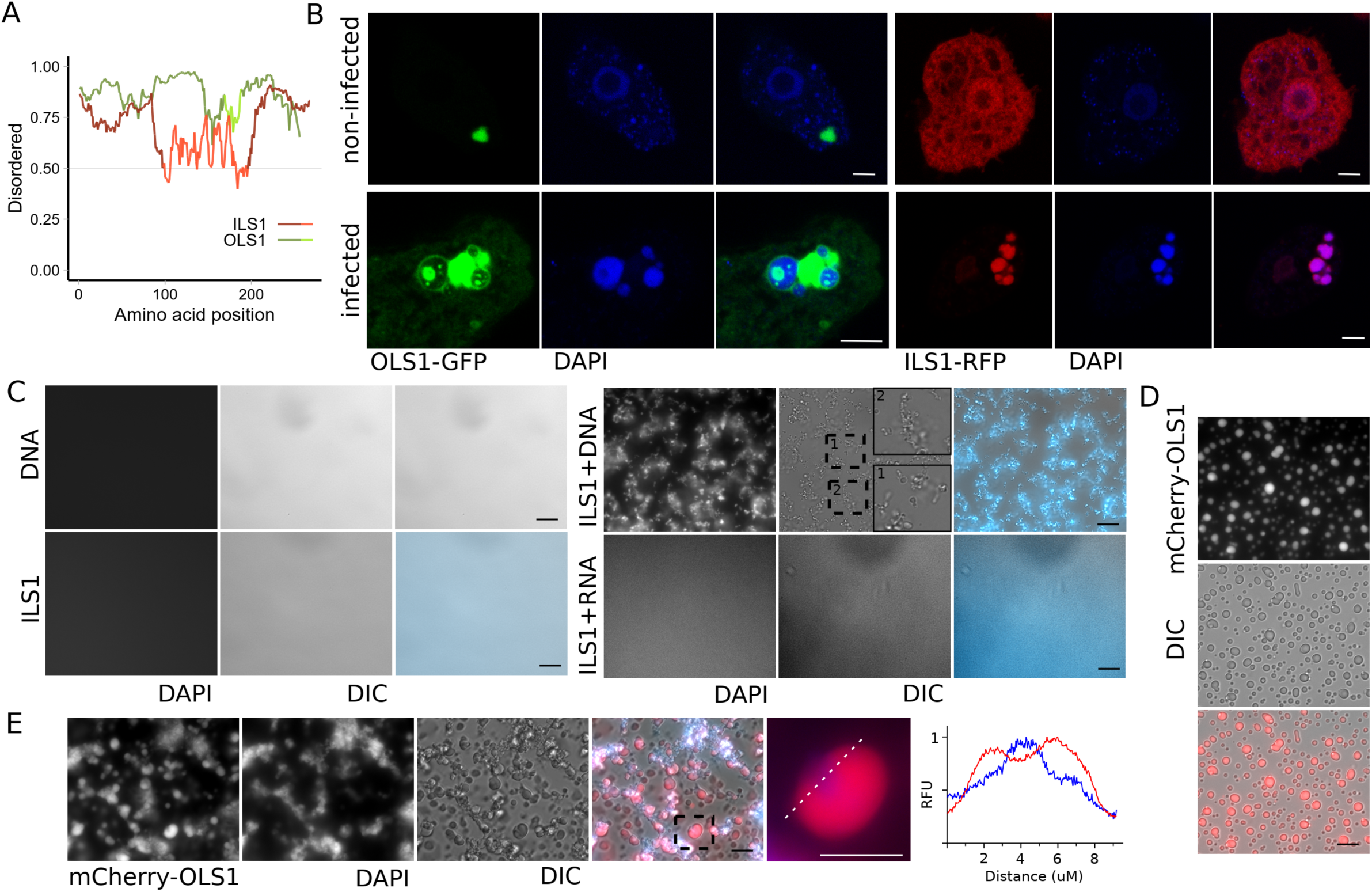
ILS1 and OLS1 are the scaffold proteins of each layer of the VF. (A) Predicted disorder tendency of OLS1 (green) and ILS1 (red).). Considered disordered regions are shown in darker color as predicted by MobiDB-lite while the numeric value was calculated by IUPred to confirm the prediction. (B) *A. castellanii* cells expressing C-terminally tagged OLS1-GFP or ILS1-RFP were infected or not with mimivirus. In absence of infection OLS1-GFP shows signs of PS at the cytoplasm of the amoeba while ILS1-RFP is shown diffused in the cytoplasm and nucleus. Upon infection, OLS1-GFP localized to the OL of the VF and ILS1-RFP to the IL of the VF. DAPI: DNA. Scale bar: 5μm. (C) *In vitro* PS of ILS1 in presence or absence of DNA or RNA. ILS1 was used at 5 µM, DNA at 10 µg/mL and RNA at 100 µg/mL. DAPI was used to confirm co-PS of protein and nucleic acid. Scale bar: 10μm. Insets were placed to allow better visualization of condensates. (D) *In vitro* PS of mCherry-OLS1 at 50mM NaCl. OLS1 was used at 5µM. Scale bar: 10μm. (E) *In vitro* PS of mCherry-OLS1 and ILS1 in presence of 10 µg/mL DNA. Both proteins were used at 5 µM. DAPI was used to confirm co-PS of protein and nucleic acid. Scale bar: 10μm. A plot profile is shown demonstrating a biphasic interaction between both phase-separated compartments.

Expression of R561 (termed Outer Layer Scaffold 1 (OLS1)) in *Acanthamoeba castellanii* demonstrated its ability to undergo PS in the amoeba cytoplasm (Figure 2B). In contrast, R252 (termed Inner Layer Scaffold 1 (ILS1)) exhibited a diffuse cytoplasmic localization (Figure 2B). Upon mimivirus infection of cells expressing both fluorescently tagged proteins, OLS1 and ILS1 re-localized to the viral factories, with OLS1 associating with the OL and ILS1 with the IL (Figure 2B). Control cells expressing only GFP or RFP did not display similar re-localizations (Figure S2A). Notably, VF client proteins, such as R336/R337, localized not only to the VF OL but also to biomolecular condensates formed by the overexpressed OLS1, strongly supporting its role as a scaffolding protein forming the VF OL (Figure S2B). Furthermore, when OLS1 and ILS1 were co-expressed, ILS1 acted as a client protein and was recruited to OLS1 biomolecular condensate (Figure S2C).

Considering that ILS1 binds DNA^30^, we reasoned that its recruitment to the VF OL would allow contact with viral DNA, enabling PS. To test this hypothesis, ILS1 was expressed and purified from *E. coli* (Figure S2D) and analyzed for PS in the presence or absence of DNA (Figure 2C). PS occurred only in the presence of DNA, as ILS1 alone remained soluble (Figure 2C). Further analysis revealed that DNA concentration significantly influenced the nature of PS (Figure S2E). At lower DNA concentrations (2.5-20 μg/mL), networks and droplets predominated, whereas higher DNA concentrations (>20 μg/mL) prompted the formation of large gels (Figure S2E). While ILS1 concentration did not affect the nature of PS (Figure S2F), it determined the speed at which it appeared. Both linear and circular DNA equally triggered PS, but mimivirus genomic DNA triggered larger gel formation with lower DNA concentration (Figure S2G). Similar results were obtained with the recombinant mCherry fused ILS1(Figure S2H). RNA did not trigger ILS1 PS (Figure 2C).

In contrast to ILS1, recombinant OLS1 (Figure S2D) made PS independently of other macromolecules (Figure 2D). PS depended on OLS1 concentrations (Figure S3A) and salt concentrations (Figure S3B). Similar to *in cellulo* conditions (Figure S2C), *in vitro* OLS1 biomolecular condensate recruited ILS1 as a client protein (Figure S3C). Finally, adding DNA to the mixture of both proteins triggered a biphasic PS, which was achieved independently of the order in which the three components were added to the mixture (Figure 2E). Altogether, OLS1, ILS1 and DNA are enough to trigger VF-like biphasic PS *in vitro*.

In order to attempt gene knockout of both genes, we generated *Acanthamoeba* transgenic lines expressing a codon-optimized version of each protein for trans-complementation^12^. Trans-complementation refers to the restoration of the function of a deleted or defective gene in a viral genome through the expression of the same gene from an independent genetic element, such as the host genome. Gene knockout of OLS1 was achieved in trans-complementing cells and clonality of recombinant viruses was demonstrated by genotyping (Figure 3A). A significant reduction of viral particle formation (approximately 100-fold reduction) was observed upon gene deletion in non-complementing cells (Figure 3B). Moreover, VF formation and/or growth was significantly inhibited, as shown by DAPI staining of infected cells (Figure 3C and 3D). This is confirmed by a lower viral DNA accumulation during infection (approximately 10-fold reduction at 6 hours post-infection) (Figure 3E). Moreover, depletion of OLS1 also reduced fusion events of VFs upon superinfection, as shown by the presence of multiple independent VFs in the same cells (Figure 3C and 3F). However, despite this strong phenotype, this gene is not essential. Thus, either the OL of the VF is composed of multiple scaffold proteins or the OL of the VF is dispensable for productive mimivirus infection. To distinguish between these two hypotheses, we analyzed the mutant infectious cycle by electron microscopy (Figure 3G). VFs of *ols1* KO viruses lack any visible OL, indicating that this compartment of the VF is dispensable for productive infection. Moreover, endogenous tagging of client proteins of the OL of the VF (like R336/R337 or R322) displayed a cytoplasmic localization upon deletion of *ols1* (Figure 3H-I and S3D-E), strongly suggesting that the role of the OL of the VF of mimivirus is to concentrate components important for VF functions at the IL or its periphery. The observed phenotypes in wild-type cells infected by *ols1* KO mutant also support the lack of additional scaffold proteins to replace OLS1.

**Figure 3.**
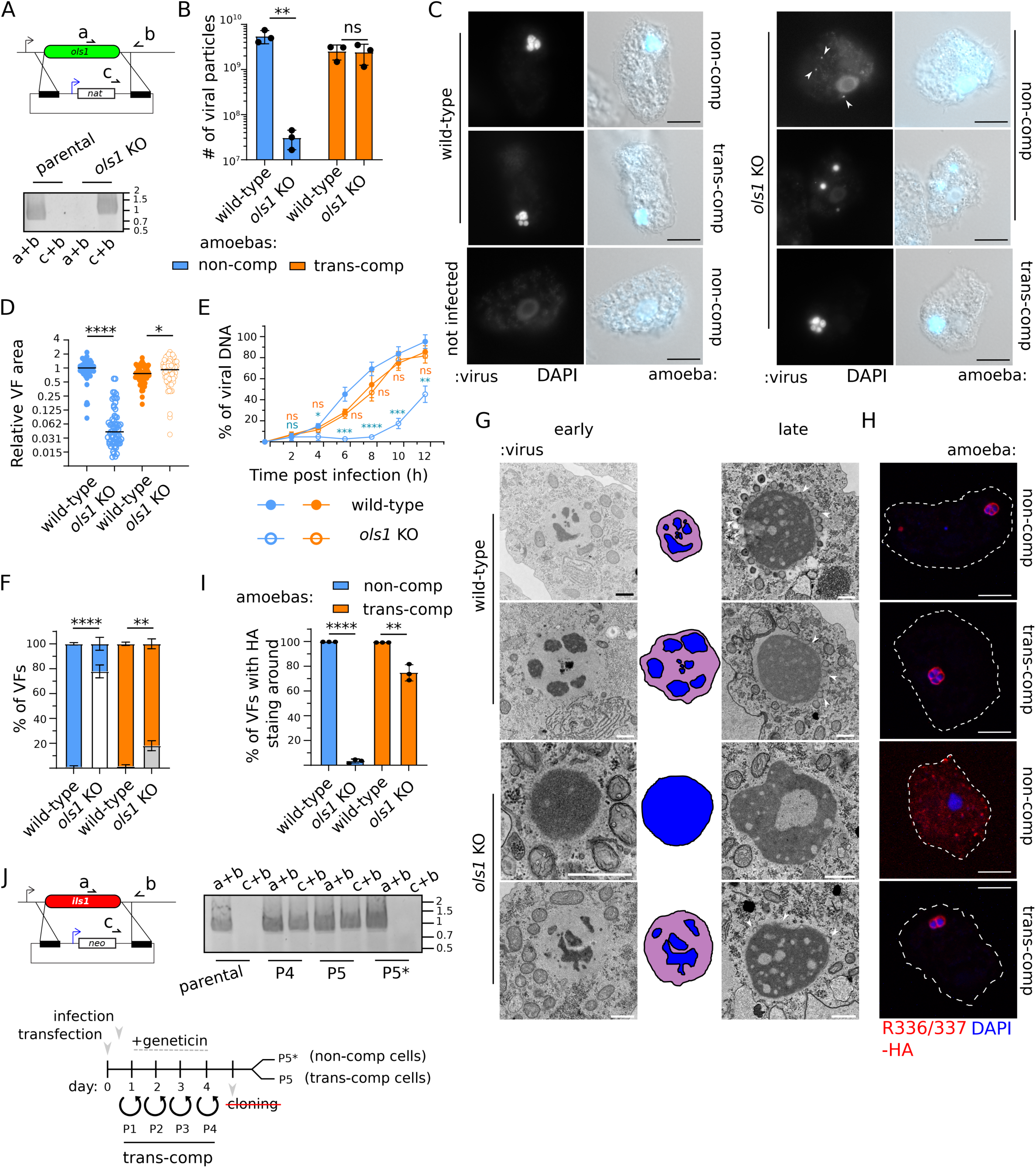
ILS1 and OLS1 play key roles in the mimivirus infection cycle. (A) Schematic representation of the vector and strategy used for *ols1* KO. Selection cassette was introduced by homologous recombination and recombinant viruses were generated, selected and cloned. *nat*: Nourseothricin N-acetyl transferase. Primers annealing locations are shown and successful KO and clonality is demonstrated by PCR. Expected size: a+b: 850bp in parental locus. c+b: 990bp in recombinant locus. (B) Quantification of the number of viruses produced upon knock-out of *ols1* in mimivirus. Infection was performed in non-complemented cells and in a trans-complementing line expressing OLS1. Data correspond to the mean ± SD of 3 independent experiments. (C) Representative light fluorescence microscopy images of *A. castellanii* cells infected with wild-type mimivirus or *ols1* KO viruses 6h pi. Infection was performed in non-complemented cells and in a trans-complementing line expressing OLS1. VFs were labelled using DAPI Scale bar: 10μm. (D) Quantification of the size of VF generated as shown in C. At least 50 VF were recorded per condition during 3 independent experiments and the area of DAPI staining was measured using ImageJ. Infection results are shown in blue or orange when generated on non-complemented cells or trans-complementing amoebae respectively. (E) DNA replication was analyzed by qPCR. Viral DNA is represented as a percentage of total DNA in the sample. Data correspond to the mean ± SD of 3 independent experiments. Infection results are shown in blue or orange when generated on non-complemented cells or trans-complementing amoebae respectively. (F) Quantification of the number of VF either fused or separated in infected cells as shown in C. At least 100 infected cells were recorded per condition. Data correspond to the mean ± SD of 3 independent experiments. Fused VFs are shown in blue or orange when generated on non-complemented or trans-complementing amoebae respectively. Separated VFs are shown in white or grey when generated on wild-type cells or trans-complementing amoebae respectively. (G) Transmission electron microscopy imaging of the mimivirus replication cycle in *A. castellanii*. Images were acquired 4-6h pi and mimivirus particles (MOI=20) were used to infect non-complemented cells or cells expressing a copy of *ols1* (trans-complementing line). Nascent virions are indicated with white arrowheads. Scale bar: 1μm. A representative cartoon indicating each of the layer of the VF is also shown: IL in blue and OL in purple. (H) Immunofluorescence demonstrating localization of client proteins from the OL of the VF. Proteins were endogenously tagged with 3xHA at the C-terminal and infection was carried out for 6h before fixation. VFs were labelled using DAPI. Scale bar: 1μm. (I) Quantification of the number of VF with clear staining of R336/337-3xHA surrounding the viral factory as shown in Figure 3H. At least 100 infected cells were quantified per condition. Data correspond to the mean ± SD of 3 independent experiments. (J) Schematic representation of the strategy used to generate the recombinant. Selection cassette was introduced by homologous recombination and recombinant viruses were generated, selected and cloning attempted as indicated by the timeline. After passage 4, viruses were split and used to infect non-complemented amoebae (P5*) or trans-complementing amoebae (P5). Populations of recombinant viruses were followed by assessing integration of the selection cassette. Expected size: a+b: 870bp in parental locus. c+b: 890bp in recombinant locus. ns (P > 0.05), * (P ≤ 0.05), ** (P ≤ 0.01), *** (P ≤ 0.001) and **** (P ≤ 0.0001).

On the other hand, we were unable to obtain clonal *ils1* KO viruses. Using the trans-complementing line, we demonstrated that *ils1* is an essential gene (Figure 3J). Future efforts will be directed at optimizing conditional depletion systems in giant viruses to study the function of essential genes by reverse genetics.

### A conserved molecular grammar allows the identification of viral factory scaffold proteins throughout the phylum *Nucleocytoviricota*

The molecular grammar of an IDR refers to compositional bias and sequence patterns in their primary structure^3,4^. This grammar allows condensation and specific recruitment of molecules, including client proteins^31–36^. We thus reasoned that due to the plethora of client proteins recruited to the VFs, despite major changes in the primary sequence of their scaffold proteins, the molecular grammar of the condensate would not easily change. Concordantly, host-expressed mimivirus (Imitervirales) OLS1-GFP and ILS1-RFP re-localized to noumeavirus (Pimascovirales) VF upon infection (Figure 4A). Moreover, host cell-expressed ILS1-RFP colocalizes with host cell-expressed C-terminal GFP-fused mel_H2B-H2A under noumeavirus-infected conditions, confirming the recruitment of ILS1-RFP to the VF of noumeavirus (Figure 4B). Thus, in order to identify scaffold proteins in all *Nucleocytoviricota*, we first predicted the “IDRome” encoded in representative genomes of isolated viruses from each viral order and extended the analysis to metagenomes from the giant virus database^37^, the permafrost^38^ and the recently discovered Egovirales^39^ (Supplementary Table 2). We then tested existing tools to classify such IDRs.

**Figure 4.**
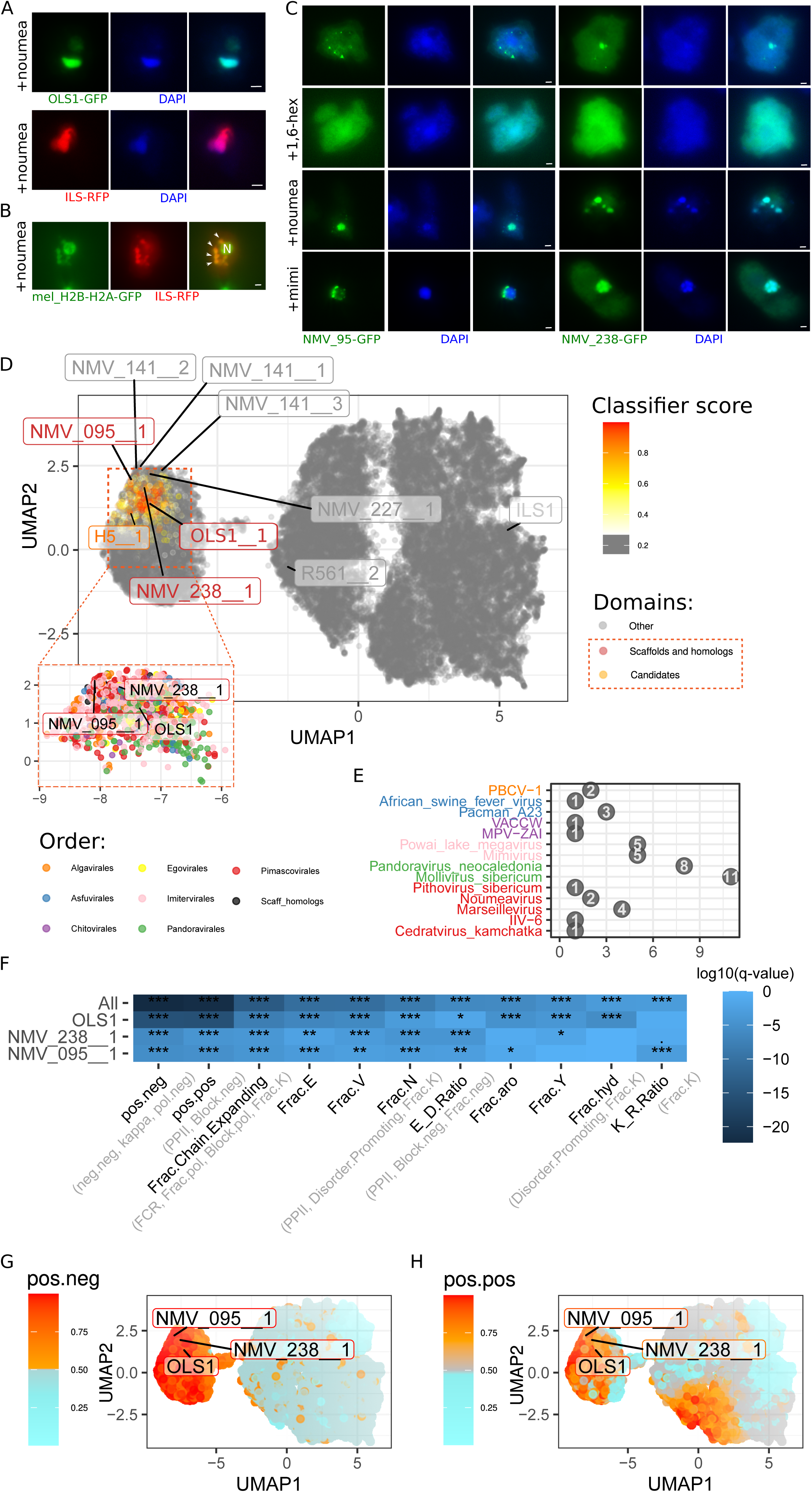
Identification of scaffold proteins by defining their molecular grammar. (A) *A. castellanii* cells expressing C-terminally tagged mimivirus OLS1-GFP or mimivirus ILS1-RFP were infected with noumeavirus. Both proteins localized to the VF of noumeavirus. DAPI: DNA. Scale bar: 1μm. (B) *A. castellanii* cells expressing C-terminally tagged melbournevirus mel_H2B-H2A-GFP and mimivirus ILS1-RFP were infected with noumeavirus. Both proteins co-localized at the VF of noumeavirus (indicated with an arrowhead). The nucleus of the host cell is marked with an N. Only mel_H2B-H2A-GFP labels the nucleus of the host cell. (C) *A. castellanii* cells expressing C-terminally tagged NMV_095 or NMV238 were either infected with noumeavirus, mimivirus or treated with 10% 1,6-hexanediol. VFs were labelled using DAPI 2-3 h pi. Scale bar: 1μm. (D) UMAP representation of the IDRs in representative genomes and metagenomics giant viruses, based on the 11 features selected for the classifier. 29,677 points are drawn, including 53 IDRs corresponding to scaffolds and homologs and 1083 candidates retrieved by the classifier (red to yellow colormap). IDRs corresponding to negative predictions are shown in grey. For visual clarity, 800 genomes of *Imitervirales* were removed because they were over-represented. Before filtering, there was 77,295 points including 3092 candidate IDRs. (E) Count of candidate proteins for phase separation predicted by the classifier in the representative genomes. (F) Final classifier features and their corrected p-value in both mimivirus and noumeavirus genomes (All) or only one of them (OLS1, NMV). Marker within the heatmap gives the level of significance: *** < 0.001, ** < 0.01, * < 0.05,. < 0.1. Features that are in parenthesis are above 0.55 correlated in the scaffold proteins and homologs. (G-H) Details of the most discriminant features on the UMAP (C). Pos.neg and pos.pos refer to the segregation of positive from negative residues or of positive residues to all other residues.

Since the IL of the VF is only present in members of the *Mimiviridae*, we excluded ILS1 from the following analysis and focused bioinformatic computations on OLS1. While tools to predict phase separation exist (ParSe v2^40^ and Molphase^41^), they predicted many candidates with the potential of achieving PS (Figure S4A). Even using a stringent threshold (0.9), molphase predicts 69% of mimivirus proteins with an IDR to do phase separation and ParSe v2, 31% of mimivirus disordered proteins. Thus, we customized specific methods to study the biochemistry of nucleolar phase separation, relying on Nardini and CIDER^36^ to increase the prediction specificity. The first IDR of OLS1 and its detectable homologues were used to determine important features separating these IDRs from the rest of the IDRome of mimivirus. Compared to the previously published method^36^, the number of scrambles for Nardini Z-score calculations was reduced (Figure S5A), and block sizes of certain amino acids were slightly adapted to allow better discrimination of OLS1 (Figure S5B). Using those 98 features (36 from Nardini and 62 from CIDER), we created machine learning classifiers that identified 4 candidate scaffold proteins in noumeavirus (Figure S4A). To experimentally challenge these predictions, the four genes were codon-optimized and expressed in the amoeba. Two showed a localization coherent with biomolecular condensates, which could be disrupted using 1,6-hexanediol *in vivo* (Figure 4C) and re-localized to the noumeavirus VF upon infection (Figure 4C). Moreover, both proteins also re-localized to the mimivirus VFs upon infection, confirming a shared molecular grammar for PS for these 2 viruses’ VFs (Figure 4C). In contrast, the other two protein candidates did not spontaneously form biomolecular condensates in the amoeba cytoplasm (Figure S4B).

We then utilized the newly confirmed noumeavirus scaffold proteins (NMV_095 and NMV_238) and detected homologs to re-train the predictive machine learning classifier. To select the features for the classifier, we compared the feature values of the three scaffold proteins IDRs and homologs to the rest of the combined IDRome of mimivirus and noumeavirus (Figure S5C-D). Final predictions of scaffold proteins in all orders of *Nucleocytoviricota* were then generated with this optimized classifier (Supplementary Table 2). Clear segregation of resulting candidate IDRs can be observed on the unsupervised UMAP representation of the features previously selected during the classifier training (Figure 4D), and putative scaffold proteins could be identified to be encoded by members of all *Nucleocytoviricota* orders (Figure 4D-E). More specifically, we identified one to five candidates from cytoplasmic viruses in all reference genomes (Figure 4E). In all these (cytoplasmic) orders, between 4.4% (*Algalvirales*) and 5.4% (*Chitovirales*) of all predicted IDRs were classified as candidate scaffold proteins for PS and VFs generation. The only exception was the Egovirales, scoring 8.7% of positive proteins in their respective IDRomes. In *Pimascovirales*, homologous proteins were identified as scaffolds in pithovirus sibericum and cedratvirus kamchatka (pv_12 and ck125, 39% identity/63% similarity), highlighting the consistency of the method. Similarly, orthopoxvirus’ H5 protein was identified as the only candidate in the vaccinia and monkeypox viruses (93% identity). On the other hand, in *Pandoravirales* (which present a nuclear DNA replication^14^ and apparently lack VFs), around 10 candidate proteins were identified. Importantly, all candidate VF scaffold IDRs predicted in all *Nucleocytoviricota* are distributed on the UMAP without order segregation, further reinforcing the shared molecular grammar for the cytoplasmic VFs (Figure 4D). As expected, scaffold IDRs have a low Anchor2 score, meaning they are predicted to stay disordered. Meanwhile, a significant portion of other IDRs have a higher Anchor2 score, indicating that they likely gain conformational order upon binding to a molecule (Figure S6A).

The most discriminative features in our classification, and thus, the features associated with the molecular grammar of the VFs, were related to charge segregation (Figure 4F-H). OLS1, NMV_095 and NMV_238 display large patches of positively and negatively charged residues (Figure S7), resulting in high Nardini positive-positive and positive-negative Z-scores. The kappa parameter^42^ is also higher for the confirmed scaffold proteins and predicted candidates (Supplementary Table 2), further pointing towards a high segregation of charges. Regardless, kappa is not part of the final classifier as the parameter is correlated to the positive-negative Nardini Z-score in the scaffold protein training set (Figure 4F). Within the negative charges, glutamate seems more important as confirmed scaffold IDRs and candidates have 15% of this residue (+/− 6), while the rest of the IDRs have 6% (+/− 8). This feature is also very important to discriminate OLS1 from ILS1 (which has only 1.4% glutamate (and other negative residues)), probably reflecting its inability to perform PS without DNA (Figure 2C). There is another group of IDRs at the center-bottom of the UMAP that scores high in positive segregation (Figure 4H). This group, for instance, resembles the candidate scaffolds in some classifier metrics but presents a low fraction of E residues (Figure S6B). As for scaffold proteins of the nucleolus, Blocks of K are higher in VF scaffold candidates than in other IDRs (0.7 +/− 0.11 vs 0), but this feature was not included in the classifier due to its high variability in scaffolds and homologs. In addition, scaffold IDRs and candidates have a relatively higher E/D ratio (0.25 +/− 0.26 vs 0 +/− 0.45) and a higher K/R ratio (0.48 +/− 0.36 vs 0.07 +/− 0.53).

Although the two first metrics are the most discriminant (Figure 4G-H), the other features are essential. They help to reduce the number of candidates from 6539 when using a classifier based only on the two first metrics to 2967 when the final classifier incorporates those 11 metrics (Supplementary Table 2). It is the integration of all these features (Figure 4G-H, Figure S6B) that makes specific discrimination possible.

### Recombinant vaccinia virus H5 and noumeavirus NMV_238 form biomolecular condensates *in vitro*

To further challenge scaffold protein predictions across the *Nucleocytoviricota*, we expressed and purified from *E. coli* the vaccinia virus H5 (as previously described ^52^) and noumeavirus NMV_238 (Figure S8A). Similar to OLS1, both H5 and NMV_238 formed biomolecular condensates in the absence of other macromolecules (Figure 5A-B). PS depended on protein concentration (Figure S8B-C) and salt concentrations (Figure S8D-E). Interestingly, in contrast to OLS1, which does not bind dsDNA, both H5 and NMV_238 were capable of binding and concentrating DNA, as shown by DAPI staining (Figure 5A-B). Moreover, condensation was slightly favored in the presence of dsDNA (Figure S8B-E). Overall, both scaffold proteins display capacities to phase separate independently of other macromolecules upon diminishing ionic strength (similar to OLS1) but display DNA binding and seclusion similarly to ILS1. Future efforts will be directed at dissecting the biophysical properties of these *in vitro*-generated biomolecular condensates.

**Figure 5.**
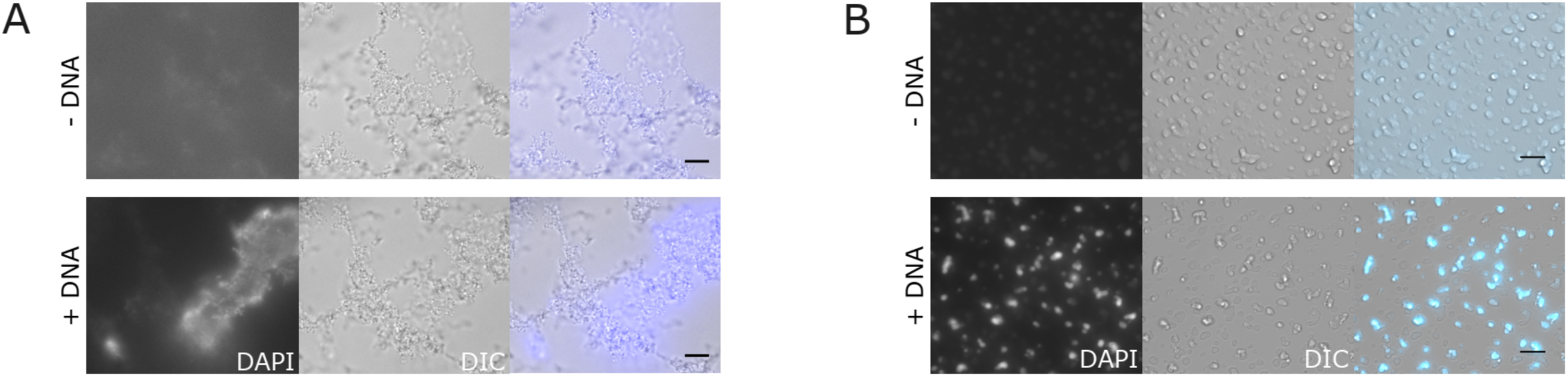
Biomolecular condensates formed by H5 and NMV_238. (A) And (B) H5 and NMV_238 form biomolecular condensates *in vitro* in the absence and presence of DNA. Phase separation was induced with 30 μM H5 (A) or 11.1 μM NMV_238 (B) in a solution containing 25 mM NaCl and 0.6 μg/mL DNA where indicated. The samples were incubated for 30 min at room temperature and stained with DAPI to confirm co-phase separation between the protein and DNA. Scale bar: 10 μm.

### Identification of client proteins demonstrates sub-compartmentalization of functions

While a previous study reported proteins tentatively localized at the VFs of mimivirus^22^, it did not differentiate proteins associated with each sub-compartments of the VFs and high rates of false positives were obtained when localization of some of these proteins was assessed by endogenous tagging (Figure S9A-B). In order to fill this gap, we performed immunoprecipitations of the two scaffold proteins and identified co-purified proteins by mass spectrometry (MS)-based proteomics (Figure 6A, Figure S9C and Supplementary Table 3). Several proteins enriched with OLS1 and/or ILS1 were endogenously tagged to confirm their localization (Figure 6B and Figure S9B). Importantly, none of the false positive examples identified from the previous proteome study were detected in these immunoprecipitations (Supplementary Table 3). Moreover, enrichment of proteins localized to the OL of the VF were co-purified with OLS1, while IL proteins or virion proteins were co-immunoprecipitated majorly with ILS1 (Figure 6A). Importantly, while this study shows a low rate of false positive identifications, other proteins not detected in the immunoprecipitations are still localized at the VF (Figure 6C). Thus, future efforts will be needed to identify the full proteome of the VFs of mimivirus.

**Figure 6.**
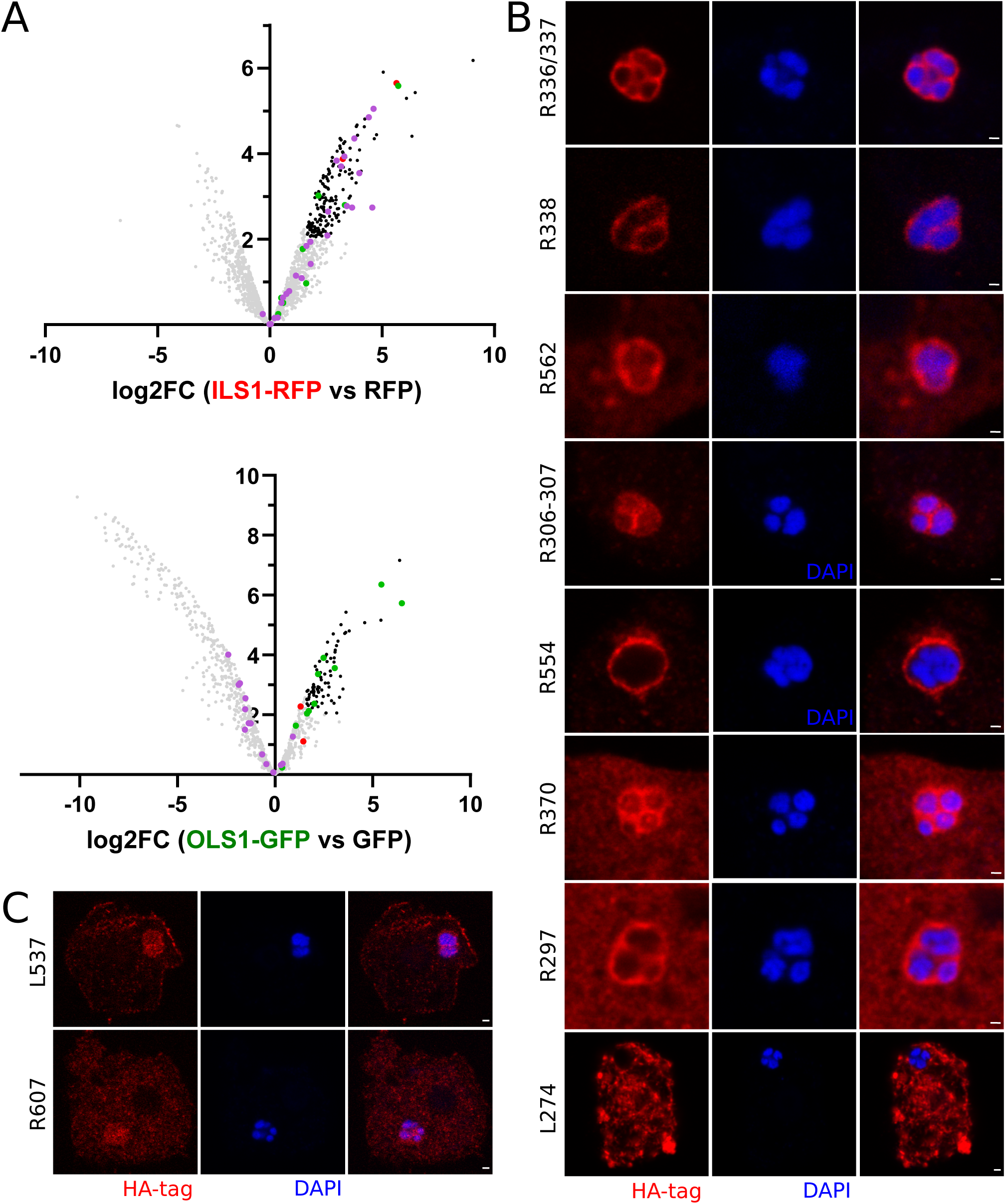
Identification of client proteins from the VF. (A) Immunoprecipitation experiments in *A. castellanii* cells expressing ILS1-RFP or OLS1-GFP and infected by mimivirus. Cells expressing RFP or GFP were utilized as controls. Crosslinking was performed prior to cell lysis. Immunoprecipitated proteins were analyzed through MS-based label-free quantitative proteomics (three replicates per condition). The volcano plots represent the −log10 (limma p-value) on y axis plotted against the log2(Fold Change bait vs control) on x axis for each quantified protein (upper panel: OLS1-GFP versus GFP, bottom panel: ILS1-RFP versus RFP). Each dot represents a protein. Proteins with log2 (FoldChange) ≥ 1.6 (FoldChange > 3) and −log10(p-value) ≥ 2 (p-value = 0.01) compare to controls, were considered significant (Benjamini-Hochberg FDR < 2%) and are shown in black. Proteins confirmed to localize at the IL, OL or virions are shown in red, green or violet respectively. Detailed data are presented in Supplementary Table 3. (B) Immunofluorescence demonstrating localization of proteins enriched in A. Proteins were endogenously tagged with 3xHA at the C-terminal of each mimivirus gene and infection was carried out for 6 hours before fixation. VFs were labelled using DAPI. Scale bar: 1μm. (C) Immunofluorescence demonstrating localization of proteins not enriched in A but still displaying a VF localization. Proteins were endogenously tagged with 3xHA at the C-terminal and infection was carried out for 6 hours before fixation. VFs were labelled using DAPI. Scale bar: 1μm.

Several host ribosomal proteins were enriched in the immunoprecipitations of OLS1-GFP (but not with ILS1-RFP), suggesting some proximity between the OL of the VF and the ribosomes (Figure 6A and Supplementary Table 3). Regardless, VFs are thought to segregate replication and transcription from translation^22^. To test the current consensus, we generated recombinant viruses or amoebae encoding tagged versions of several proteins involved in the central dogma (Figure 7A-D and Figure S9A). As previously suggested^22^, host ribosomes (Figure 7C and Figure S10A) and the viral translation-associated protein SUI1 (Figure 7D) were excluded from the VFs. On the other hand, the virally encoded eIF4e localized both at the cytoplasm of the host and the OL of the VF (Figure 7D). In eukaryotes, besides its classical cytoplasmic function in translation initiation, eIF4E also localizes at the nucleus, where it participates in the export of a subset of mRNA^43^. If such export function is conserved in the virally encoded eIF4E remains to be explored, but it would explain the dual localization of the protein.

**Figure 7.**
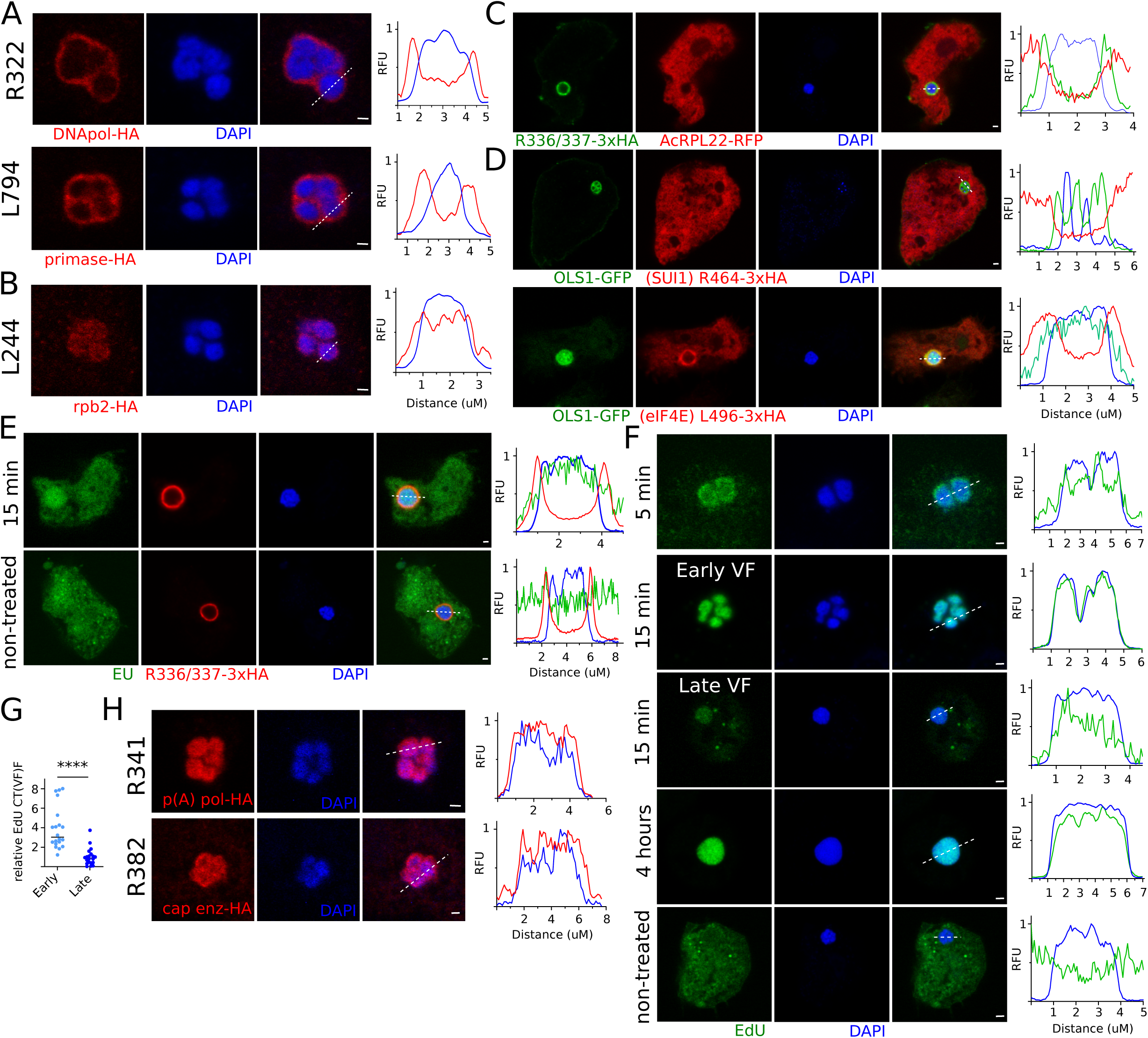
Sub-cellular and sub-organelle compartmentalization of the proteins involved in the central dogma. (A) Immunofluorescence demonstrating localization of proteins associated to DNA replication. Proteins were endogenously tagged with 3xHA at the C-terminal and infection was carried out for 6 hours before fixation. VFs were labelled using DAPI. Line profiles (bottom) corresponding to the white dashed line show fluorescence patterns. Scale bar: 1μm. (B) Immunofluorescence demonstrating localization of the RNA polymerase subunit 2 (rpb2), associated to transcription. Protein was endogenously tagged with 3xHA at the C-terminal and infection was carried out for 6 hours before fixation. VFs were labelled using DAPI. Line profiles (bottom) corresponding to the white dashed line show fluorescence patterns. Scale bar: 1μm. (C) Immunofluorescence demonstrating localization of host ribosomal protein RPL22-RFP, associated to translation. Protein is expressed from a second copy plasmid encoding *rpl22-rfp* and infection was carried out for 6 hours before fixation. IL and OL of the VFs were labelled using DAPI and R366/337-3xHA respectively. Line profiles (bottom) corresponding to the white dashed line show fluorescence patterns. Scale bar: 1μm. (D) Immunofluorescence demonstrating localization of viral proteins associated to translation. Proteins were endogenously tagged with 3xHA at the C-terminal and infection was carried out for 6 hours before fixation. IL and OL of the VFs were labelled using DAPI and OLS1-GFP respectively. Line profiles (bottom) corresponding to the white dashed line show fluorescence patterns. Scale bar: 1μm. (E) Detection of RNA synthesis by EU labelling. Viral infection by wild type viruses was allowed to proceed for 4-6 hours and labelling time is indicated. IL and OL of the VFs were labelled using DAPI and R366/337-3xHA respectively. Scale bar: 1μm. (F) Detection of DNA synthesis by EdU labelling. Viral infection by *thymidylate synthase* KO viruses was allowed to proceed for 4-6 hours and labelling time is indicated. IL of the VFs was labelled using DAPI. Scale bar: 1μm. (G) Quantification of the corrected total VF fluorescence (CT(VF)F) of EdU labeling as shown in F. 20 VFs were recorder during 3 independent experiments and the intensity of EdU staining was measured using ImageJ. ns (P > 0.05), * (P ≤ 0.05), ** (P ≤ 0.01), *** (P ≤ 0.001) and **** (P ≤ 0.0001). (H) Immunofluorescence demonstrating localization of proteins associated to mRNA maturation. Proteins were endogenously tagged with 3xHA at the C-terminal and infection was carried out for 6 hours before fixation. VFs were labelled using DAPI. Line profiles (bottom) corresponding to the white dashed line show fluorescence patterns. Scale bar: 1μm.

On the other hand, proteins associated with replication and transcription are incorporated in the VFs but sub-compartmentalized. While replication proteins localized to the OL of the VF, transcription proteins accumulated at the IL (Figure 6A and Figure 7A-B). These data might indicate that replication and transcription are occurring in different compartments of the VF. To test that hypothesis, pulse-labeling of mRNAs using 5-ethynyluridine (EU) for 15 minutes strongly suggests that mRNA production site is located at the IL of the VF (Figure 7E), colocalizing with the RNA polymerase (Figure 7B). On the other hand, we were unable to efficiently label DNA by 5-Ethynyl-2-deoxyuridine (EdU) in wild type viruses (Figure S10B). We reasoned that due to the high AT richness of mimivirus genome^44^, *de novo* synthesis of dTMP would be particularly efficient in infected cells, allowing thymidine to outcompete EdU for its incorporation into the DNA (Figure S10C). Thus, we generated knockout recombinant viruses on the virally encoded thymidylate synthase (TS), concordantly blocking viral *de novo* synthesis of dTMP (Figure S10D-E). EdU labeling significantly improved in these viruses, allowing for the visualization of DNA replication with pulses of EdU labelling as short as 5 minutes (Figure 7F and Figure S10F). Similarly to what was observed in the vaccinia virus^45^, pulse labeling of DNA replication in late VF resulted in a lower fluorescence intensity than labelling in early VF (Figure 7F-G). These data strongly indicate that DNA replication decreases during the late stages of infection. Moreover, 5-minute labelling with EdU showed an enrichment of newly synthesized DNA at the periphery of the IL of the VF (Figure 7F). Concordantly, DNA replication likely occurs at the interface between the IL and the OL, where DNA and the DNA polymerase get in contact. Finally, mRNA processing (including capping and Poly(A) synthesis) localized at the IL of the VF, indicating that maturation of pre-mRNA occurs at the same sub-compartment as transcription (Figure 7H). Overall, a significant sub-compartmentalization of functions is observed at mimivirus VFs.

## Discussion

The molecular grammar of an IDR of a scaffold protein determines the nature of the interaction to achieve PS and the selective recruitment of client proteins^31–36,46^. Here, we demonstrate that members of the *Nucleocytoviricota* phylum share a common molecular grammar which allows the formation of their VFs. Thus, the common ancestor of all *Nucleocytoviricota* already possessed a VF with this molecular grammar. Such conservation is likely driven by the complex number of client proteins recruited to the VFs, which would need to change their biochemical/biophysical properties simultaneously if the molecular grammar of the VF suddenly changes. Such a scenario is parsimoniously unlikely. Regardless, it is unclear if all scaffold protein IDRs share a common origin and diversification occurred by shuffling protein fragments^47^ or if the same molecular grammar emerged in multiple IDRs on different occasions by convergent evolution^48^. Which advantages the virus gains by modifying the scaffold proteins that form their VFs remains to be addressed but might be associated with emergent traits of different VFs (including the appearance of an IL in mimivirus or the endoplasmic reticulum (ER) wrapping in poxviruses^49^). Moreover, the condensates formed *in vitro* by OLS1, ILS1, H5 and NMV_238 showed striking differences, ranging from individualized and spherical (OLS1) to more network-like condensates (H5 and ILS1). This visual evidence suggests different biophysical properties of the condensates. Future efforts will be directed to dissect the differences within the context of the multiple post-translational modifications these proteins suffer during infection (*i.e.* H5^50^). Importantly, such grammar allowed us to identify previously neglected but well-characterized scaffold proteins in other members of the phylum. In *Poxviridae*, H5 was predicted as the sole candidate for a scaffold protein for PS. H5 is an essential protein for the infectious cycle of the vaccinia virus^51^ and was coined as a hub protein due to its importance in DNA replication, transcription and virion morphogenesis^52^. All those phenotypes correlate with a scaffolding function for PS. Moreover, H5 binds DNA^52^ (a function modulated by phosphorylation^50^) and localizes as puncta upon heterologous expression in mammalian cells (indicating spontaneous PS of the protein)^51^. Interestingly, when vaccinia virus uncoating occurs, early viral proteins associated with DNA replication localize to cytoplasmic puncta, including H5^53,54^. We propose that these puncta (known as prereplication foci) are likely formed by PS using H5 as a scaffold protein. Upon maturation of the prereplication foci, VF gets surrounded by the ER membranes^49^. Regardless, H5 strongly accumulates inside the VF^54,55^, and the VFs dissolve in the presence of 1,6-hexanediol. Moreover, VFs of the vaccinia virus have previously been shown to coalesce when present in the same cell^56^. These data support the idea that PS is a major driver of the VF formation regardless of the ER wrapping. Moreover, it has been proposed that H5 would be a key component for VF enlargement and wrapping by the ER^53^. Overall, previously published data on H5 strongly supports its role as an unrecognized scaffold protein for PS. In *Marseilleviridae*, at least two scaffold proteins localize to the VFs with differential transcriptional expression patterns^57^. This allows us to hypothesize that NMV_095 might initiate VF formation while NMV_238 would allow its expansion and maturation. Further experiments will be needed to corroborate this hypothesis. In *Pandoravirales*, multiple IDRs containing a similar molecular grammar to VF scaffold proteins were identified. Regardless, these viruses do not form VFs and transfer their DNA into the nucleus of their host. In evolutionary terms, it is parsimonious to assume that a fully cytoplasmic infectious cycle style of the majority of the members of the phylum originated prior to the nuclear one^58^, as the origin of *Nucleocytoviricota* predated the origin of eukaryotes (and thus, the nucleus)^59^. Thus, during the transition from cytoplasmic to nuclear viruses, the transfer of the DNA into the nucleus would generate a VF, which is no longer needed to achieve replication and transcription but would still keep the functions associated with virion morphogenesis. Nonetheless, further experiments would be needed to characterize such “Virion Factories” potentially assembled by nuclear giant viruses.

Mimivirus VFs are highly compartmentalized organelle-like structures (Figure 8). Biogenesis of the VFs starts by utilizing the cores as nucleating points. Similar observations were previously made on vaccinia virus, indicating some conserved mechanisms of nucleation^55^. Mimivirus VFs then develop into multilayered structures that contain at least two distinctive phases. The OL of the VF, formed by OLS1, acts as a selective barrier and recruits VF proteins. Importantly, while we hypothesize that the OL of mimivirus VFs is the phylogenetically conserved phase between different *Nucleocytoviricota*, OLS1 is dispensable in mimivirus. We theorize that the presence of the IL allows the protection of the genome of the virus in the absence of the OL and, despite losing the ability to selectively recruit proteins to the VF and considering the permissive conditions of the laboratory, the IL is sufficient to achieve successful infection. The DNA replication machinery localizes to the OL of the VF and maximizes DNA replication only at the interphase between OL and IL. This interphase is also the site of the assembly of virions, and the competition between these two processes might at least partially explain the decrease in DNA synthesis during the late stages of the infection cycle. The IL of the VF contains proteins associated with transcription, which are also packaged into the virions to establish a new cycle of infection (including the RNA polymerase, transcription factor, RNA processing, etc.). Thus, the IL seems to be a compartment analogous to the internal content of the virion core, which is sufficient to restart RNA transcription of early genes upon infection of a new cell^15^. Moreover, ILS1 has recently been proposed to work on mimivirus DNA condensation for its incorporation in the viral particle^30^. Finally, translation occurs outside of the VF, a feature differentiating mimivirus from vaccinia virus^60^. Importantly, how mRNAs are exported from the VFs is still unknown. Regardless, since neither ILS1 nor OLS1 requires RNA for PS, a simple model where RNA is not retained by the IL or OL of the VF can be envisioned. In such a case, diffusion would be sufficient to deliver mRNAs into the cytoplasm of the infected cell.

**Figure 8.**
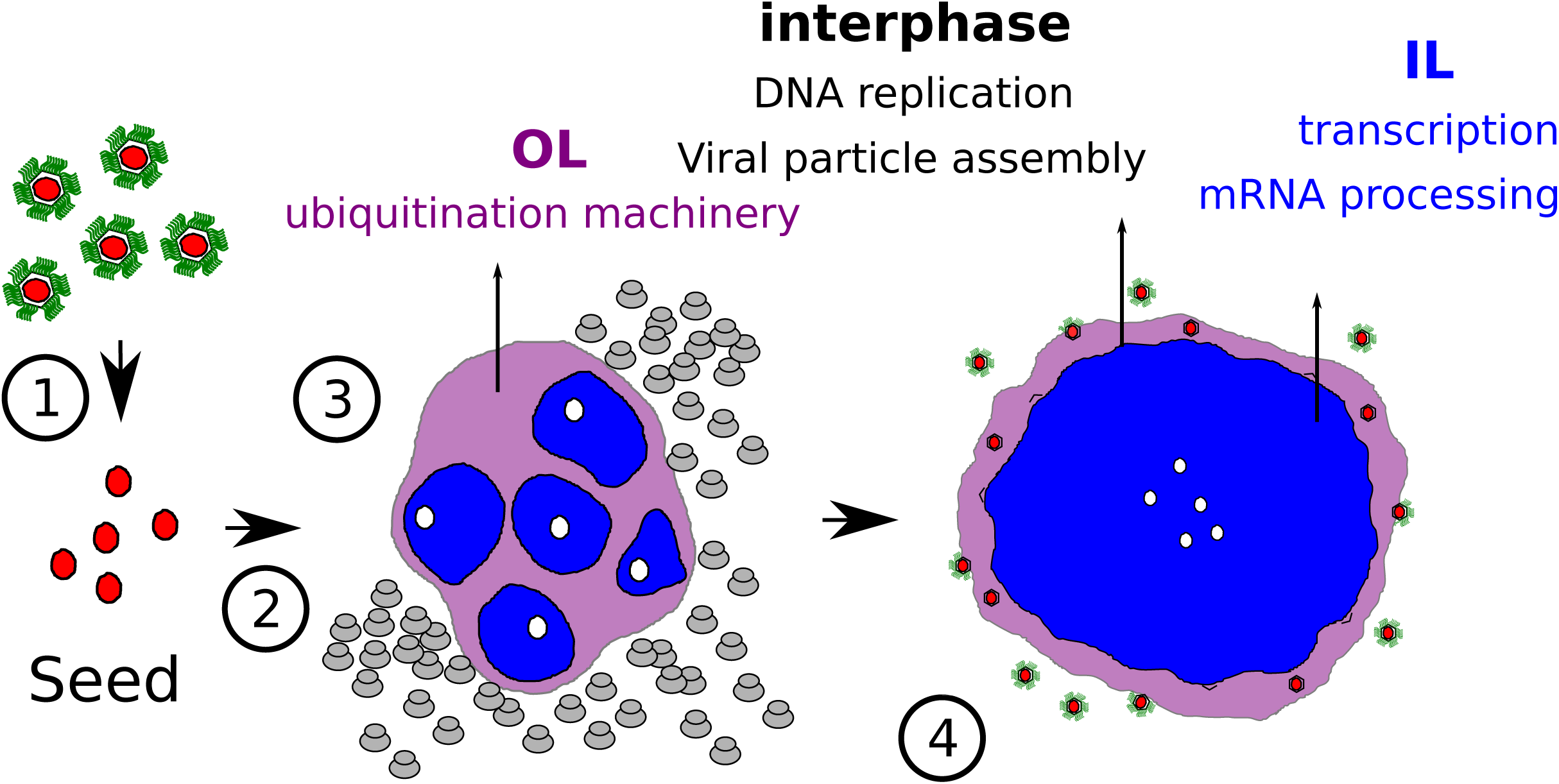
Schematic model of mimivirus infection. Schematic depiction of the proposed model of mimivirus VF. Infective mimivirus (1) are internalized to the cell by phagocytosis and deliver an internal structure (core, red) containing the viral DNA (2). Individual viral factories are nucleated around the internalized cores (white). VFs are then generated by multilayered phase separation. Outer layer (OL, purple) quickly fuse in case of superinfection of host cells while Inner layer (IL, blue) fuse later. OL concentrates proteins associated to post-transcriptional regulation like ubiquitination while IL concentrates DNA as well as transcription and mRNA maturation. DNA synthesis and virion morphogenesis occur at the interface between OL and IL while glycosylation of virions (green) occurs upon exit of the VF. Early VFs (3) and late VFs (4) are depicted.

Overall, *Nucleocytoviricota* VFs are biomolecular condensates with a shared phylogenetic history and a common molecular grammar. Such discovery raises new questions, including how cytoplasmic viruses interact upon infection of the same host cell (can VFs from different viruses fuse?), how biomolecular condensates accommodate spatiotemporally of the different functions required for VFs’ multiple roles and open the door to the development of generalist drugs to inhibit *Nucleocytiviricota* viral infections.

## Author contributions

S.R., conceptualization, methodology, validation, formal analysis, investigation, data curation, writing (original draft, review and editing), software, visualization; A.S., conceptualization, methodology, validation, formal analysis, investigation, data curation, writing (original draft, review and editing), software, visualization; A.B.M, conceptualization, methodology, validation, formal analysis, investigation, data curation, writing (original draft, review and editing), software, visualization; A.L., investigation and data analysis; L.D., investigation and data analysis; C.G., investigation and data analysis; F.T., investigation and data analysis; L.B., investigation and data analysis; N.O-D., investigation, data analysis and writing (review & editing); Y.C., supervision, methodology, validation, formal analysis and writing (review & editing); M.B., investigation, data analysis, supervision and writing (review & editing); M.L., supervision and writing (review & editing); S.J., supervision and writing (review & editing); C.A., conceptualization, supervision, writing (review & editing) and funding acquisition; H.B., conceptualization, methodology, validation, formal analysis, investigation, data curation, visualization, writing (original draft, review and editing), supervision, project administration and funding acquisition.

## Acknowledgements

The authors thank Matias Estaras Hermosel for critical discussion to improve the manuscript. The authors also thank Christian Betzel, Gamaleldin I. Harisa, Chunfu Zheng, Anouk Willemsen, Andrea Maria Guarino, Yves Gaudin, Juliana Cortines and Kekungu-u Puro for the comments raised on the preprint version of the manuscript which were taken into consideration for the revision. We thank the members of the PiCSL-FBI core facility (Nicolas Brouilly, Fabrice Richard and Aïcha Aouane), IBDM, AMU-Marseille, and on the IMM imaging platform (Artemis Kosta and Hugo Le Guenno). The proteomic experiments were partially supported by Agence Nationale de la Recherche under projects ProFI (Proteomics French Infrastructure, ANR-10-INBS-08) and GRAL, a program from the Chemistry Biology Health (CBH) Graduate School of University Grenoble Alpes (ANR-17-EURE-0003). NO and MB gratefully acknowledge the Advanced Bioimaging Unit at the Institut Pasteur Montevideo & Universidad de la República for their support and assistance in the present work. Funded by the European Union (grant agreement No. 101160452, ERC-2024-STG to H.B., and grant agreement No 832601 ERC-2018-ADG to C.A). Views and opinions expressed are, however, those of the authors only and do not necessarily reflect those of the European Union or the European Research Council (ERC). Neither the European Union nor the granting authority can be held responsible for them.

## Methods

### *A. castellanii* growth and virus production

The following viral strains have been used in this study: Acanthamoeba polyphaga mimivirus^44^, noumeavirus^58^, pithovirus sibericum^61^, pacmanvirus lost city^62^, mollivirus sibericum^27^. Ten infected 75 cm² tissue-culture flasks were plated with fresh *Acanthamoeba* cells for virus production. After lysis completion, the cultures were recovered and centrifuged for 5 min at 500 × *g* to remove the cellular debris. The virus was pelleted by a 45 min centrifugation at 6,800 × g prior to purification. The viral pellet was then resuspended and washed twice in PBS, layered on a discontinuous CsCl gradient (1.2/1.3/1.4/1.5 g/cm3), and centrifuged at 100,000 × g overnight. An extended protocol is shown in ^63^.

#### Vaccinia virus

The Modified Vaccinia virus Ankara (MVA) strain was propagated in BHK-21 cells. The viral stock was generated in 25 cm² tissue-culture flasks with cell monolayers at a multiplicity of infection (MOI) of 0.1. After a 1-hour adsorption period, the inoculum was removed, and the cells were incubated at 37 °C in a humidified atmosphere with 5% CO₂. The cultures were collected when 80-90% of the cell monolayer exhibited cytopathic effects (CPE), typically 2-3 days post-infection. The cultures were centrifuged at 1500 x *g* for 5 minutes, after which the supernatant was aliquoted and stored at −80 °C. The viral titer was determined using a TCID50 endpoint dilution assay.

For experiments involving 1,6-hexanediol treatment, viral infections were carried out in Vero cells using an MOI of 10.

Both BHK-21 and Vero cells were cultured in Dulbecco’s Modified Eagle’s Medium (DMEM) supplemented with 10% fetal bovine serum (FBS) and an antibiotic-antimycotic solution (penicillin 100 units/mL, streptomycin 100 μg/mL, and amphotericin B 0.25 μg/mL). During viral infections, the FBS concentration was adjusted to 1%.

*Acanthamoeba castellanii* (Douglas) Neff (American Type Culture Collection 30010TM) cells were cultured at 32 °C in 2% (wt/vol) proteose peptone, 0.1% yeast extract, 100μM glucose, 4mM MgSO_4_, 0.4mM CaCl_2_, 50 μM Fe(NH_4_)_2_(SO_4_)_2_, 2.5 mM Na_2_HPO_4_, 2.5 mM KH_2_PO_4_, pH 6.5 (home-made PPYG) medium supplemented with antibiotics [Ampicillin 100 μg/mL, and Kanamycin 25 μg/mL]. 100 μg/mL Geneticin G418 or Nourseothricin was added when necessary.

### Generation of DNA constructs

#### Vectors for endogenous tagging

vAS1 plasmid was utilized for endogenous tagging^9^. 500 bp homology arms were introduced at the 5’ and 3’ end of the selection cassette to induce homologous recombination with the viral DNA. Each cloning step was performed using the Phusion Taq polymerase (ThermoFisher) and InFusion (Takara). Before transfection, plasmids were digested with ApaI/EcoRI/HindIII and NotI. Primers utilized are shown in Supplementary Table 4.

#### Second copy vectors for expression in A. castellanii

The encoding genes for OLS1 (R561), ILS1 (R252), NMV_95, NMV_141, NMV_227, and NMV_238 were codon optimized for amoeba expression and amplified by PCR to be cloned into different amoeba expression vectors (PAM1, PAM2, PAM3, PAM10, or Vc241^9^). The plasmid was linearized by NdeI, and the gene was inserted using InFusion Takara. The primers utilized are shown in Supplementary Table 4.

#### Vectors for gene knockout of ols1 (r561) and ils1 (r252)

vHB47 was used as the plasmid for gene knock-out^9,10^. 500 bp homology arms were introduced at the 5’ and 3’ end of the selection cassette to induce homologous recombination with the viral DNA. Each cloning step was performed using the Phusion Taq polymerase (ThermoFisher) and InFusion (Takara). Before transfection, plasmids were digested with ApaI/EcoRI/HindIII and NotI. Primers utilized are shown in Supplementary Table 4.

#### Vector for gene knockout of thymidylate synthase

vAS1 was used as the plasmid for gene knock-out^9,10^. 500 bp homology arms were introduced at the 5’ and 3’ end of the selection cassette to induce homologous recombination with the viral DNA. Each cloning step was performed using the Phusion Taq polymerase (ThermoFisher) and InFusion (Takara). Before transfection, plasmids were digested with HindIII and NotI. Primers utilized are shown in Supplementary Table 4.

### Establishment of viral lines

#### Generation of recombinant viruses

Recombinant viruses were generated as described step by step in ^9^. Briefly, 1.5×10^5^ *Acanthamoeba castellanii* cells were transfected with 6 μg of linearized plasmid using Polyfect (QIAGEN) in phosphate saline buffer (PBS). One hour after transfection, PBS was replaced with PPYG and cells were infected with mimivirus for 1 hour with sequential washes to remove extracellular virions. 24 hours after infection, the new generation of viruses (P0) was collected to infect new cells. An aliquot of P0 viruses was utilized for genotyping to confirm the integration of the selection cassette. A new infection was allowed to proceed for 1 hour, then washed to remove extracellular virions and nourseothricin and/or geneticin was added to the media. Viral growth was allowed to proceed for 24 hours. This procedure was repeated again before removing the nourseothricin and/or geneticin selection to allow recombinant viruses to expand more rapidly. Once the viral infection was visible, the selection procedure was repeated one more time. Viruses produced after this new selection round were used for genotyping and cloning. The selection utilized for each virus generation is indicated in the “Generation of DNA constructs” section.

#### Cloning and genotyping

Cloning and genotyping of recombinant viruses are extensively described in ^9^. Briefly, 50,000 *A. castellanii* cells were seeded on 6-well plates with 2 mL of PPYG. After adhesion, viruses were added to the well at a multiplicity of infection (MOI) of 2. One hour post-infection, the well was washed 5 times with 1mL of PPYG, and cells were recovered by scraping the well. Amoebae were then diluted until obtaining a suspension of 1 amoeba/μL. One μL of such suspension was added in each well of a 96-well plate containing 1×10^3^ uninfected *A. castellanii* cells and 200 μL of PPYG. Wells were later monitored for cell death, and 100 μL collected for genotyping. Genotyping was performed using Terra PCR Direct Polymerase Mix (Takara) following manufacturer specifications. Primers utilized for each genotyping are detailed in Supplementary Table 4.

### Protein expression and purification

Protein expression and purification were performed as previously described^30,52^. Briefly, cultures were grown in Luria-Bertani (LB) medium containing ampicillin until an optical density of 0.5-0.6 (600nm). Bacterial expression was then induced with 0.3 mM isopropyl-1-thio-β-d-galactopyranoside (IPTG), and bacteria were incubated at 16°C for 15 h with constant shaking. Cells were then collected and centrifuged at 4,000 x *g* prior to resuspension in 50 mM Tris pH 7.4, 5 mM imidazole, 1 M NaCl, 5% glycerol, complete anti-protease tablets (Roche). Cells were lysed by sonication before clearing by centrifugation at 16,000g for 30 min. The soluble fraction was loaded on a Hi-Trap Chelating HP 1 ml pre-packed column (GE Healthcare/Cytiva). Elution was carried out in 50 mM Tris pH 7.4, 5 mM imidazole, 500mM NaCl, 5% glycerol. Protein fractions obtained were pooled and later dialyzed with 20 mM Tris pH 7.5, 100 mM NaCl and 5% glycerol. Purified proteins were concentrated using Amicon® Ultra 15 ml centrifugal filters and stored in aliquots at −80°C. Protein concentrations were determined using the nanodrop and their theoretical epsilon.

### Phase separation assays

All droplet formation assays were performed in the absence of crowding agents. Proteins were diluted into specified buffers in a final assay volume of 100 μL. When indicated, RNA or DNA was added to the mixture. Samples were visualized on a non-binding 96-well microplate (Greiner bio-one). DNA used for most of the experiments corresponds to circular PAM8 (5138 bp). PAM8 was linearized using NdeI restriction enzyme when specified. The genomic DNA of mimivirus was utilized when specified (approx. 1.2Mbp). The RNA used was extracted from mimivirus infected cells using the RNeasy midi kit (Qiagen) according to the instructions provided by the manufacturer.

### Immunofluorescence and fluorescence microscopy

*A. castellanii* cells were grown on poly-L-lysine coated coverslips in a 12-well plate, infected or not with viruses and fixed with PBS containing 3.7% formaldehyde for 20 min at room temperature. When required, immunofluorescence was performed as described step by step in ^9^. After three washes with PBS buffer, coverslips were mounted on a glass slide with 4 μl of VECTASHIELD mounting medium with DAPI and the fluorescence was observed using a Zeiss Axio Observer Z1 inverted microscope using a 63x objective lens associated with a 1.6x Optovar for DIC, mRFP or GFP fluorescence recording.

Vero cells were grown on poly-L-lysine treated 96-well plate, infected or not with viruses and fixed with 4% formaldehyde in PBS (Fixative Solution, Invitrogen) with DAPI for 15 min at room temperature. The fluorescence was observed using a 40x objective lens in an Olympus IX81 inverted microscope using the µManager software.

### 1,6-hexanediol treatment

To disrupt viral factories, 1,6-hexanediol (240117, Sigma-Aldrich) was diluted in cell culture media at 10% w/v as previously described^2^. Cell culture media were replaced with media containing 10% 1,6-hexanediol or fresh culture media and incubated for 10 min at 32 or 37 °C before fixation.

### Virion production quantification

Optical density was utilized for viral quantification as previously described^9^. The purity of the viral samples was analyzed by microscopy^63^ or genotyping^9^, as previously described.

### DNA quantification

Viral genomes or gDNA from infected amoebae were purified using the Wizard genomic DNA purification kit (PROMEGA). To determine the amplification kinetic, the fluorescence of the EvaGreen dye incorporated into the PCR product was measured at the end of each cycle using the SoFast EvaGreen Supermix 2× kit (Bio-Rad, France). For each experiment, a standard curve using gDNA of purified viruses was performed in parallel. For each point, a technical triplicate was performed. The primers utilized are shown in Supplementary Table 4.

### Electron microscopy imaging

Extracellular virions or *A. castellanii*-infected cell cultures were fixed by adding an equal volume of PBS with 2% glutaraldehyde and 20 min incubation at room temperature. Cells were recovered and pelleted 20 min at 5,000 × g. The pellet was resuspended in 1 mL PBS with 1% glutaraldehyde, incubated at least 1 h at 4 °C, and washed twice in PBS prior to coating in agarose and embedding in Epon resin. Each pellet was mixed with 2% low melting agarose and centrifuged to obtain small flanges of approximately 1mm^3^ containing the sample coated with agarose. These samples were then prepared using the osmium-thiocarbohydrazide-osmium method: 1 h fixation in 2% osmium tetroxide with 1.5% potassium ferrocyanide, 20 min in 1% thiocarbohydrazide, 30 min in 2% osmium tetroxide, overnight incubation in 1% uranyl acetate, 30 min in lead aspartate, dehydration in increasing ethanol concentrations (50, 70, 90 and 100% ethanol) and embedding in Epon-812. Ultrathin sections of 70 nm were observed using a FEI Tecnai G2 operating at 200 kV^58^.

### Immunoprecipitation

At 6 hours post-infection *Acanthamoeba castellanii* infected cells were harvested, washed in PBS and crosslinked. Crosslinking was performed using 1% formaldehyde for 10 minutes, followed by quenching in 0.125M glycine solution in PBS. Cells were washed once and lysed in co-immunoprecipitation buffer (0.2% v/v Triton X-100, 50 mM Tris-HCl, pH 8, 150 mM NaCl) in a protease inhibitor cocktail (Roche). Cells were sonicated on ice and centrifuged at 14,000 r.p.m. for 30 min at 4°C. Supernatants were then immunoprecipitated using anti-HA, anti-GFP, or anti-RFP antibodies, as previously described^64^.

### MS-based proteomic analyses

Proteins eluted from co-IP experiments were either separated by SDS-PAGE (HA-tagged proteins and WT control, one replicate per condition) or stacked (GFP- and RFP-tagged proteins and respective controls, three replicates per condition) in the top of a 4-12% NuPAGE gel (Invitrogen) before Coomassie blue staining and in-gel digestion using modified trypsin (Promega, sequencing grade) as previously described^65^. For co-IP experiments with HA-tagged proteins and WT control, the bands corresponding to large chains of immunoglobulins were prepared and analysed separately from the rest of the samples. The resulting peptides were analyzed by online nanoliquid chromatography coupled to MS/MS (Ultimate 3000 RSLCnano and Q-Exactive HF, Thermo Fisher Scientific) using gradients of 140 min, 35 min or 80 min for the eluates of HA-tagged proteins and of the WT control, the bands corresponding to the immunoglobulin large chains in HA-co-IPs, and GFP- and RFP-tagged proteins and the corresponding controls, respectively. The MS and MS/MS data were acquired using Xcalibur (Thermo Fisher Scientific). The mass spectrometry proteomics data have been deposited to the ProteomeXchange Consortium via the PRIDE^66^ partner repository with the dataset identifier PXD054803.

Peptides and proteins were identified by Mascot (version 2.8.3, Matrix Science) through concomitant searches against the following homemade databases: *A. castellanii* nuclear genome (17’625 sequences), *A. castellanii* mitochondrial genome (40 sequences), mimivirus (979 sequences), and contaminants classically found in proteomic analyses (keratins, trypsin… 250 sequences). Trypsin/P was chosen as the enzyme and two missed cleavages were allowed. Precursor and fragment mass error tolerances were set at respectively at 10 and 20 ppm. Peptide modifications allowed during the search were: Carbamidomethyl (C, fixed), Acetyl (Protein N-term, variable) and Oxidation (M, variable). The Proline software^67^ (version 2.3) was used for the compilation, grouping and filtering of the results (conservation of rank 1 peptides, peptide length ≥ 6 amino acids, false discovery rate of peptide-spectrum-match identifications < 1%^68^, and minimum of one specific peptide per identified protein group). Proline was then used to perform a MS1-based label-free quantification of the identified protein groups based on specific and razor peptides.

Statistical analysis was performed using the ProStaR software^69^. Proteins identified in the contaminant database were discarded. For the dataset of HA-tagged proteins, after log2 transformation, abundance values were normalized using median centering. To be considered enriched with a HA-tagged bait protein, a protein must show a normalized abundance at least four times higher in the bait protein eluate than in the WT control eluate and be identified with a minimum of three spectral counts. For the dataset of GFP- and RFP-tagged proteins, proteins detected in less than three replicates of one condition were discarded. After log2 transformation, abundance values were normalized using the variance stabilizing normalization (vsn) method, before missing value imputation (SLSA algorithm for partially observed values in the condition and DetQuantile algorithm for totally absent values in the condition). Statistical testing was conducted with limma, whereby differentially expressed proteins were selected using a log2 (Fold Change) cut-off of 1.6 and a p-value cut-off of 0.01, allowing to reach false discovery rates inferior to 2% according to the Benjamini-Hochberg estimator. Proteins detected in fewer than three replicates in the condition in which they were most abundant were manually invalidated (p value = 1).

### EU and EdU labelling

*Acanthamoeba* cells were grown on glass coverslips and infected with mimivirus at MOI of 10. Incorporation and visualization of EU or EdU was performed as previously described^15^ utilizing Click-iT™ EdU Cell Proliferation Kit for Imaging, Alexa Fluor™ 488 dye and Click-iT™ RNA Alexa Fluor™ 488 Imaging Kit Invitrogen. Briefly, after labeling for the specified time with 100 μM EdU or 1 mM EU, cells were fixed with 3.7% paraformaldehyde for 20 min, washed, permeabilized with 0.5% Triton X-100, and incubated with the Click-iT™ reaction mixture as indicated by the manufacturer.

### Statistics and reproducibility

All data are presented as the mean ± s.d. of 3 independent biological replicates (n = 3), unless otherwise stated in the figure. All data analyses were carried out using Graphpad Prism. The null hypothesis (α = 0.05) was tested using unpaired two-tailed Student’s t-tests.

### Database constitution with the molecular grammar of IDRs

A database was constituted with genomes of isolated viruses: African swine fever virus (GCA_003815255.1), cedratvirus kamchatka (GCA 031200085.1), Invertebrate iridescent virus 6 (IIV-6, GCA 000838105.1), marseillevirus (GCF 001806195.1), mimivirus (GCA 000888735.1), mollivirus sibericum (GCF 001292995.1), monkeypox virus Zaire (MPV-ZAI, GCA 000857045.1), noumeavirus (GCF 002005685.1), pandoravirus neocaledonia (GCF 003233915.1), paramecium bursaria chlorella virus 1 (PBCV-1, GCA 000847045.1), pithovirus sibericum (GCA 000916835.1), powai lake megavirus (GCA 002924545.1), vaccinia virus WR (VACCW, GCA 900236015.1). Predicted proteins from the Giant virus database^37^ from the 8 large genomes from permafrost metagenomics (PRJEB47746), and from *Egovirales*^39^ were also included. Intrinsically disordered regions in all proteins were predicted using MobiDB-lite v3.10.0^70^.

Each IDR’s molecular grammar was extracted by estimating residue binary patterns and physicochemical properties

For the residue binary patterns, the python package Nardini v1.1.1 was used. It infers Z-scores for all IDRs based on the positive, negative, polar, hydrophobic, aromatic residues and alanines, prolines or glycines^4^. To infer the Z-scores, it performed 50,000 scrambles, meaning the sequences are shuffled in order to calculate a Z-score of the real value within the artificial values. The optimal number of scrambles was chosen by comparing Z-scores from 10 to 500,000 scrambles to the Z-scores obtained with 1,000,000 scrambles using the IDRs of mimivirus and homologs of its scaffold proteins.

Compositional data and physical and chemical properties of IDRs were predicted with localCIDER v0.1.21^71^ as in ^36^ with the addition of the kappa estimation. The block lengths of certain residues were also counted similarly to King et al.^36^, only counting the residues of interest (no mismatch) and subtracting opposite residues (positives vs negatives, hydrophobic vs polar).

The resulting Nardini and CIDER features were then normalized by subtracting the median and dividing by the inter-quantile range previously calculated on all IDRs, including from metagenomes. All Z-scores and normalized features were then saturated by the sigmoid function^72^.

### Constitution of the reference IDR and homologs database

In order to prepare the training of the classifier, the full set of IDRs has been split in two: a positive set, corresponding to experimentally confirmed scaffold proteins and homologs, and a negative set consisting of all the other IDRs found in the previously described database.

Homologs of OLS1, NMV_095 and NMV_238 were recovered in two steps. First, the proteins were aligned to the Giant virus database and the permafrost metagenomes^38^ by MMseqs2 v.12^73^ with an e-value cutoff of 1e-5. Secondly, sequences were aligned with t-coffee v13.41.0^74^ and HMM models were constructed with HMMER v3.3.2^75^ and searched for in the metagenomic database. Sequences were considered as homologs only for alignments with e-values <1e^−10^. The homologous proteins were then aligned again with t-coffee. Only IDRs aligned to the reference (OLS1 1, NMV_095 1 or NMV_238 1) with at least 10 overlapping amino acids were kept. Important features were determined in the same way as for the final classifier: wilcoxon signed-rank test from the scipy package v. 1.13.1 were performed to compare reference IDRs and homologs to the rest of the IDRs of mimivirus or noumeavirus. The p-values were corrected by the false_discovery_control function. Only features with a significantly lower variance, given by variance comparison and a levene test, were considered to further ensure that we compared a homogeneous population. For all the significantly relevant features, the Euclidean distance was calculated and IDRs with a distance above a manually set threshold were discarded after inspection of heatmaps presenting those features for each IDR. Three IDRs were then manually removed from the homologs of NMV_238 as they were too distant from the scaffolds in the UMAP. The final homologous IDRs used for the training set are given in Table S7.

### Prediction of the VF scaffold proteins

Several classifiers were developed with two primary goals: first, to identify candidate proteins in noumeavirus that could correspond to OLS1, and second, to predict OLS1 proteins in other genomes using the generalized knowledge gained from validating noumeavirus candidates.

To set up an SVM-based predictor, we calculated the qvalues of all IDR features differentiating the positive dataset (OLS1 and homologs, then NMV_095 and NMV_238 and homologs when confirmed) from the negative dataset. Features were sorted on their q-value and classifiers were created based on the first N features (N varying from 2 to 20). This higher limit was set to ensure the number of features did not exceed approximately 1/10th the number of IDRs in the training set, reducing the risk of overfitting (Fig.S5D). Then, to minimize redundancy and ensure the selection of distinct features, the Spearman correlation in the scaffold proteins (reference IDR and homologs) was used to identify and exclude overly correlated features (Figure S5C).

We tested SVM classifiers with both linear and radial basis function (rbf) kernels, with a C parameter of 5 and a gamma parameter of 5. Data points were weighted according to the following scheme, aiming for the lowest possible false positive and false negative rates, even if the negative set is originally highly populated: for instance, the weights of all negative IDRs sum up to 20, the weights of all scaffold and homologs IDR to 10, and the weights of the 3 experimentally confirmed IDRs sum up to 6. Finally, experimentally rejected noumeavirus candidates NMV_141 and NMV_227 were given an extra weight of 1 and 2, respectively, in the negative set to ensure the classifier adequately reflects their experimental rejection.

The final SVM classifier was built on an rbf kernel considering 11 features whose Spearman correlation coefficient is under 0.55. These features included two Nardini features (positive-negative Z-score, positive-positive Z-score), and 9 CIDER features (proportion of chain expanding residues, fraction of E, V, N residues, E versus D ratio, fraction of aromatic, Y, and hydrophobic residues, and K versus R ratio).

For comparison we also tested MolPhase^41^ and ParSe v2^40^, two tools that predict phase separation scaffold proteins.

Figures were drawn on R v. 4.2.1 (https://www.R-project.org/) and the ggplot2 package^76^ UMAPs were drawn with the uwot package (https://doi.org/10.48550/arXiv.1802.03426)

### Data and code availability

All the codes used for the bioinformatic analysis and for the VFCpredict tool developed in this study are available at https://src.koda.cnrs.fr/igs/vfcpredict.

## Glossary

PS: phase separation
VF: viral factories
IL: inner layer
ILS1: inner layer scaffold 1
OL: outer layer
OLS1: outer layer scaffold 1
IDR: intrinsically disordered regions.

## Supplemental Figures

**Supplementary Figure 1.**

(A) Transmission electron microscopy imaging of the mimivirus VFs formed in the cytoplasm of *A. castellanii*. Potential coalescent events of the inner layer are highlighted by arrowheads. Image was acquired 6h pi at a MOI=20. A cartoon representing the two layers of the viral factory is also shown. Inner layer (IL) is shown in blue while outer layer (OL) is shown in purple.

(B) Quantification of the experiments is shown in Figure 1D and Figure 1H. Data correspond to the mean ± SD of 3 independent experiments. Quantification performed based on DAPI staining is shown in blue, while quantification performed based on mollivirus MCP-RFP in red. Light blue or orange bars represent quantification performed in the presence of 10% 1,6-hexanediol treatment. ns (P > 0.05), * (P ≤ 0.05), ** (P ≤ 0.01), *** (P ≤ 0.001) and **** (P ≤ 0.0001).

(C) Cells expressing OLS1-GFP or mel_H2B-H2A-GFP were infected with mimivirus or noumeavirus, respectively. After treatment with mock or 1,6-hexanediol, cells were collected, counted and processed for western blot analysis. Lysates produced from an equal number of cells were loaded, and quantifications were performed and normalized.

(D) Cells infected with mimivirus or noumeavirus were collected after treatment with mock or 1,6-hexanediol and counted. DNA was extracted as described in Materials and Methods. qPCRs were performed as described in Materials and Methods, and viral DNA content was normalized by the total DNA extracted.

(E) Representative light fluorescence microscopy images of *A. castellanii* cells infected with different viruses belonging to *Nucleocytoviricota*. VFs were labelled using DAPI and treatment with hypotonic media (5% PYG in water) was performed for 15 minutes after pithovirus (6 hpi) and pacmanvirus (6 hpi) infection. Scale bar: 1μm.

(F) Quantification of the experiments shown in Figure 1D and Figure S1E. Data correspond to the mean ± SD of 3 independent experiments. Quantification performed based on DAPI staining is shown in blue. Light blue bars represent quantification performed in presence of 5% PYG in water treatment. ns (P > 0.05), * (P ≤ 0.05), ** (P ≤ 0.01), *** (P ≤ 0.001) and **** (P ≤ 0.0001).

(G) Live-cell imaging of mimivirus infected-*A. castellanii* expressing OLS1-GFP as a marker of the OL of the VF. Infection was allowed to proceed for 3 h and recording was performed every 2 s. Two viral factories (marked with magenta and red arrowheads) juxtaposed during the recording and drifted together for the entire recording time (1 hour and 30 minutes). Scale bar: 10μm.

(H) Silver stain and western blot associated to the immunoprecipitation performed with 3xHA tagged R562, R505 and R336/R337.

**Supplementary Figure 2.**

(A) *A. castellanii* cells expressing GFP or RFP were infected or not with mimivirus. In the absence of infection, GFP and RFP are diffused in the cytoplasm and nucleus. Upon infection, no major changes in localization are detected. DAPI: DNA. Scale bar: 5 μm.

(B) Immunofluorescence demonstrating localization of client protein R336/R337-3xHA to the OL of the VF and OLS1-GFP biomolecular condensate. White arrowhead shows the VF. Unfilled arrowhead indicate the biomolecular condensate formed by the excess of OLS1. Scale bar: 5 μm.

(C) *A. castellanii* cells expressing C-terminally tagged OLS1-GFP and ILS1-RFP illustrating that ILS1 is a client protein of OLS1. DAPI: DNA. Scale bar: 5 μm.

(D) Purified ILS1, OLS1, mCherry-ILS1 and mCherry-OLS1 proteins analysis on Sodium Dodecyl Sulphate-Polyacrylamide gel (SDS-PAGE). The mCherry fusion displayed significant contaminations with *E. coli* proteins but all results were confirmed with the non-tagged proteins. Loading order: mcherry-ILS1 (MW 69.5), mcherry-OLS1 (MW 70.2), V5-ILS1 (MW 30.8) and V5-OLS1 (MW 31.5). Proteins migrates at a higher molecular weight than predicted.

(E) *In vitro* PS of ILS1 in presence of DNA. ILS1 was used at 5 µM and DNA ranged from 0 to 80 µg/mL. DAPI was used to confirm co-PS between protein and nucleic acid. Scale bar: 10 μm.

(F) *In vitro* PS of ILS1 in presence of DNA. DNA was used at 10 µg/mL and ILS1 ranging from 0 to 8 µM. DAPI was used to confirm co-PS between protein and nucleic acid. Scale bar: 10 μm.

(G) *In vitro* PS of ILS1 in presence of DNA. ILS1 was used at 5 µM and DNA at 10 µg/mL. Circular and linear plasmid as well as mimivirus genomic DNA were compared. DAPI was used to confirm co-PS between protein and nucleic acid. Scale bar: 10 μm.

(H) *In vitro* PS of mCherry-ILS1 in presence of DNA. ILS1 was used at 5 µM and DNA at 40 µg/mL. Circular and linear plasmid as well as mimivirus genomic DNA were compared. DAPI was used to confirm co-PS between protein and nucleic acid. Scale bar: 10 μm.

**Supplementary Figure 3.**

(A) *In vitro* PS of mCherry-OLS1 at 50mM NaCl. mCherry-OLS1 was used at different concentrations. Scale bar: 10μm.

(B) *In vitro* PS of mCherry-OLS1 at different NaCl concentrations. mCherry-OLS1 was used at 5 µM. Scale bar: 10μm.

(C) *In vitro* PS of OLS1 at 50mM NaCl. OLS1 was used at 5 µM. Different concentration of mCherry-ILS1 were added to the mix. Scale bar: 10μm.

(D) Immunofluorescence demonstrating localization of client proteins from the OL of the VF. Proteins were endogenously tagged with 3xHA at the C-terminal, and infection was carried out for 6 hours before fixation. VFs were labelled using DAPI. Scale bar: 1μm.

(E) Schematic representation of the vector and knock-in (KI) strategy utilized for endogenous tagging of *r322* and *r336/337*. Selection cassette was introduced by homologous recombination, and recombinant viruses were generated, selected and cloned. *nat*: Nourseothricin N-acetyl transferase. Primer annealing locations are shown, and successful KI as clonality is demonstrated by PCR. Expected sizes are indicated in the figure.

**Supplementary Figure 4.** Exploration of different methods to predict scaffold proteins in noumeavirus with OLS1 and homologs.

(A) Correlation matrix between different classifiers positive proteins predictions. The white number in each cell indicates the number of shared predictions between two classifiers. The cell color corresponds to the fraction of shared predictions (the number of shared predictions normalized by the number of positive proteins in the classifier identifying the lowest number of positive proteins). Numbers in black give the number of representative genomes in which the two classifiers predicted at least one scaffold protein. The 4 noumeavirus proteins predicted as positive by at least one of our methods are shown in the graph under the heatmap in grey, red, orange, and blue, respectively. Each line corresponds to a link between the protein and its associated classifier.

(B) *A. castellanii* cells expressing C-terminally tagged NMV_141 or NMV_227. Scale bar: 1μm.

**Supplementary Figure 5.** Improvement of feature calculations and optimization of classifiers

(A) Optimal number of bootstraps performed by Nardini to estimate a Z-score. The error is given as the absolute difference with the reference Z-score obtained using 1 million sequence randomizations of mimivirus IDRs. Only Nardini features with an inter-quantile range above 0 are considered.

(B) Modification of the block calculation method from King *et al*.^36^. The wilcoxon p-values for the reference IDR and homologs against the rest of mimivirus IDRs were compared. Note: To ensure that the features used in this figure were shared and relevant, we only kept the ones with significantly lower variance in the predicted scaffold proteins compared to the negative class.

(C) F1-score of classifiers with OLS1, NMV_238 and NMV_095 and homologs as training sets compared to the correlation threshold above, which features can be clustered together. The final classifier clusters feature a Spearman correlation coefficient above 0.55, as correlation thresholds of 0.55 and higher all provide high f1-scores.

(D) F1-score variations with respect to the number of features considered by the classifier. The final classifier is based on selected 11 uncorrelated features, as the f1-scores do not improve when using more uncorrelated features.

**Supplementary Figure 6.** Distribution of classifier features and control features across the UMAP representation of IDRs of *Nucleocytoviricota*.

(A) Anchor2 values across the UMAP. Red points are for possible truly disordered IDRs having at least 15 amino acids with an Anchor2 score under 0.5.

(B) Final classifier features across the UMAP representation of IDRs based on those 19 features plus the positive-positive and positive-negative Nardini Z-score shown in Figure 4, highlighting each feature contribution to the classifier and in validated scaffold proteins. ‘Frac.’ refers to the fraction, and the chain-expanding residues include E (glutamate), D (aspartate), R (arginine), K (lysine), and P (proline). ‘PPII’ denotes the propensity to adopt polyproline II conformations. Additionally, ‘pol’ represents polar residues, while ‘hyd’ corresponds to hydrophobic residues.

**Supplementary Figure 7.** Charge segregation in reference scaffold proteins. The positive and negative charge distribution given by CIDER is shown as red for positive and blue for negative. The grey-shaded rectangle highlights the position of the IDRs. These 3 identified scaffold proteins show similar alternating positive and negative patches across the IDR. NCPR: Net charge per residue.

**Supplementary Figure 8.**

(A) Purification of H5 and NMV_238 proteins by His-trap nickel-affinity column. *Left:* Coomassie-stained SDS-PAGE gel of H5 purification. *Right:* Coomassie-stained SDS-PAGE gel of NMV_238 purification. The molecular weight (MW) ladder is indicated on the left of the gel. Lanes 1, total protein; Lane 2, soluble protein; Lane 3, pellet; Lane 4, flowthrough; Lane 5, wash with 25 mM Imidazole; Lane 6, wash with 50 mM Imidazole; Lane 7, elution with 250 mM Imidazole.

(B) H5 and (C) NMV_238 *in vitro* phase separation at different protein concentrations in the absence or presence of DNA. Phase separation observed at increasing protein concentrations with 25 mM NaCl and 0.6 μg/mL of DNA (right panel). Scale bar: 10 μm.

(D) H5 and (E) NMV_238 *in vitro* phase separation in different salt (NaCl) concentrations in the absence or presence of DNA. Phase separation observed at increasing NaCl concentrations with 30 μM H5 (D) or 11.1 μM NMV_238 (E) and 0.6 μg/mL of DNA (right panel). Scale bar: 10 μm.

**Supplementary Figure 9.**

(A) Immunofluorescence demonstrating the localization of proteins identified at the VF in ^22^ that could not be confirmed by endogenous tagging. Proteins were endogenously tagged with 3xHA at the C-terminal, and infection was carried out for 6 hours before fixation. VFs were labelled using DAPI. Scale bar: 2μm.

(B) Schematic representation of the vector and knock-in (KI) strategy utilized for endogenous tagging of client proteins. The selection cassette was introduced by homologous recombination, and recombinant viruses were generated, selected and cloned. *nat*: Nourseothricin N-acetyl transferase. Primer annealing locations are shown, and successful KI as clonality is demonstrated by PCR. Expected sizes are indicated in the figure.

(C) Silver stain gels and western blots corresponding to the immunoprecipitation performed with GFP or RFP tagged OLS1 or ILS1, respectively.

**Supplementary Figure 10.**

(A) *A. castellanii* cells expressing RPL22-RFP. RPL22-RFP was detected in the nucleolus and the cytoplasm of non-infected cells. DAPI: DNA. Scale bar: 2 μm.

(B) Detection of DNA synthesis by EdU labelling. Viral infection by wild type viruses was allowed to proceed for 4-6 hours and labelling time is indicated. IL of the VFs was labelled using DAPI. Scale bar: 2 μm.

(C) Cartoon representing the two pathways for incorporation of deoxy-thymidine into the DNA. TK: Thymidylate kinase. TS: Thymidylate Synthase.

(D) Cartoon representing the strategy to disrupt the Thymidylate Synthase gene without disrupting the N-terminal Dihydrofolate reductase domain.

(E) Efficient disruption of *thymidylate synthase* of mimivirus and clonality of recombinant viruses was demonstrated by PCR. Primer annealing locations are shown in Figure S6C. Expected sizes are indicated in the figure.

(F) Quantification of the relative number of VFs with EdU labeling in wild-type viruses or *thymidylate synthase* KO is shown. Data correspond to the mean ± SD of 3 independent experiments. At least 100 VFs were counted during each experiment. ns (P > 0.05), * (P ≤ 0.05), ** (P ≤ 0.01), *** (P ≤ 0.001) and **** (P ≤ 0.0001).

**Supplementary Table 1. MS-based characterization of R505, R562 and R336-R337 interactomes.**

**Supplementary Table 2. Prediction of the IDRome and putative scaffold proteins throughout the *Nucleocytoviricota*.**

**Supplementary Table 3. MS-based quantitative proteomic identification of OLS1(R561)-GFP and ILS1(R252)-RFP binding partners.**

**Supplementary Table 4. Primers used in this study.**

